# Inhibition of pyrimidine biosynthesis targets protein translation in AML

**DOI:** 10.1101/2021.09.24.461627

**Authors:** Joan So, Alexander C. Lewis, Lorey K. Smith, Kym Stanley, Lizzy Pijpers, Pilar Dominguez, Simon J. Hogg, Stephin J. Vervoort, Ricky W. Johnstone, Gabrielle McDonald, Danielle B. Ulanet, Josh Murtie, Emily Gruber, Lev M. Kats

**Affiliations:** The Peter MacCallum Cancer Centre, Melbourne, 3000 VIC, Australia; The Sir Peter MacCallum Department of Oncology, University of Melbourne, Parkville, 3052 VIC, Australia; Human Oncology and Pathogenesis Program, Memorial Sloan Kettering Cancer Center, New York, New York, USA; Servier Pharmaceuticals, Boston, MA, USA

## Abstract

The mitochondrial enzyme dihydroorotate dehydrogenase (DHODH) catalyzes one of the rate-limiting steps in *de novo* pyrimidine biosynthesis, a pathway that provides essential metabolic precursors for nucleic acids, glycoproteins and phospholipids. DHODH inhibitors (DHODHi) are clinically used for autoimmune diseases and are emerging as a novel class of anti-cancer agents, especially in acute myeloid leukemia (AML) where pyrimidine starvation was recently shown to reverse the characteristic differentiation block in AML cells. Herein we show that DHODH blockade rapidly shuts down protein translation in leukemic stem cells (LSCs) by down-regulation of the multi-functional transcription factor YY1, has potent activity against AML *in vivo* and is well tolerated with minimal impact on normal blood development. Moreover, we find that ablation of CDK5, a gene that is recurrently deleted in AML and related disorders, increases the sensitivity of AML cells to DHODHi. Our studies provide important molecular insights and identify a potential biomarker for an emerging strategy to target AML.

## Main

AML is an aggressive malignancy with few effective treatment options and extremely poor outcomes in the majority of cases. The 5-year overall survival rate is less than 30%, and for a large proportion of patients that have unfavorable prognostic factors the median survival is less than one year ^1^. Hence, there is an urgent unmet need to develop novel therapeutic strategies, particularly those that engage different mechanisms of action compared with drugs in current clinical use ^2^.

Pyrimidine bases are components of many biological macromolecules including DNA and RNA, and are essential for cell growth. Although mammalian cells can acquire pyrimidines from salvage pathways, most cells rely predominantly on *de novo* synthesis to meet their metabolic requirements. DHODH is a flavoprotein that is localized on the inner-mitochondrial membrane and catalyzes the fourth step of *de novo* pyrimidine biosynthesis, the ubiquinone-mediated oxidation of dihydroorotate to orotate. As an enzyme that is associated with the electron transport chain, DHODH links nucleotide synthesis with mitochondrial bioenergetics and ROS production ^3–5^.

Inhibition of DHODH has been proposed as a therapeutic strategy in a range of human diseases from viral infection, to auto-immunity and cancer ^3–5^. The FDA-approved DHODHi leflunomide has been clinically utilized as an immune-suppressant for the treatment of rheumatoid arthritis and multiple sclerosis, with its efficacy proposed to stem from selective anti-proliferative effects on high-affinity T-cells ^6^. Leflunomide and another DHODHi, brequinar, have been trialed in various cancers and although occasional durable responses were observed, the initial data was insufficient to support further clinical development ^3–5^. Notably however, recent findings suggesting selective efficacy of DHODHi in hematological malignancies have prompted renewed interest in DHODH as an anti-cancer target and have spurred extensive efforts to develop more potent and selective next-generation inhibitors ^7–11^. Intriguingly, AML cells have been reported to undergo myeloid maturation in response to pyrimidine starvation^7–10^, suggesting a link between nutrient availability and cell fate that can be exploited as a form of “differentiation therapy”.

To facilitate further development and clinical deployment of DHODHi, there remains a need for a comprehensive analysis of molecular and cellular responses, especially in rare leukemia stem cells (LSCs) that underpin disease progression and therapy resistance ^12^. Herein we used AG636, a novel potent DHODHi that has been extensively validated by biophysical studies and metabolic profiling^11^, to characterize the link between nucleotide metabolism and therapeutic efficacy. Our findings demonstrate that DHODH inhibition has anti-proliferative and pro-differentiation activity *in vivo* and potent activity against multiple AML sub-types. By integrating data across a panel of syngeneic mouse models and human AML cells lines, we characterize molecular pathways that are directly triggered by DHODHi and identify a novel predictive biomarker and combination strategy.

## Results

### DHODH inhibition has potent anti-leukemic activity in MLL-rearranged AML *in vivo*

Inhibition of DHODH has been reported to induce differentiation of AML blasts *in vitro* and *in vivo*, but the pathways that underpin this phenotypic response remain unknown ^7–10^. To gain insights into the mode of action of DHODH-targeting drugs we focused initially on an aggressive chemo-refractory syngeneic murine AML model driven by doxycycline-inducible expression of MLL-AF9 and constitutive expression of oncogenic Nras^G12D^ (hereafter referred to as MN)(Extended Data Fig. 1a)^13,14^. We and others have previously shown that leukemic cells are addicted to continued expression of the MLL-AF9 oncoprotein and that silencing it drives terminal myeloid differentiation ^14,15^. We reasoned that comparing MLL-AF9 depletion with AG636 treatment may aid in distinguishing primary cellular and molecular changes triggered directly by DHODH inhibition from secondary changes that occur as a consequence of differentiation.

We first performed a longitudinal study monitoring disease progression and differentiation status of leukemic cells at weekly intervals by flow cytometric analysis of peripheral blood. We administered AG636 at a dose of 100mg/kg body weight b.i.d. on days one to five of a seven-day cycle, a regimen that was well-tolerated (Extended Data Fig. 1b). AG636 marginally reduced disease progression after one cycle of treatment and we observed evidence of differentiation as indicated by reduced expression of cKit and increased expression of Ly6G (Fig. 1a-b). Excitingly, following cycle two we observed disease regression in the AG636 group, and four cycles of treatment was sufficient to induce clinical complete remission with no detectable MN cells in the peripheral blood of 10 out of 12 mice (Fig. 1b-c). In the majority of animals these responses were sustained for more than eight weeks following drug withdrawal and extensive FACS analysis of the bone marrow at the endpoint failed to detect any minimal residual disease in 9 survivors. Two mice did relapse however (M#15 and M#16) and one other animal required euthanasia although no tumor cells were detected in its blood or bone marrow (Extended Data Fig. 1c). To determine whether relapsed MN cells were resistant to DHODH inhibition, we transplanted these cells into a cohort of secondary recipients. We also transplanted a separate control cohort with MN cells from a vehicle-treated animal (M#30) that had never been exposed to AG636. In both cohorts, drug treatment (using a reduced two-week regimen) was equally effective, demonstrating that even though leukemic cells could occasionally persist during four weeks of therapy, they retained a high degree of AG636 sensitivity at relapse (Extended Data Fig. 1d).

**Fig. 1:**
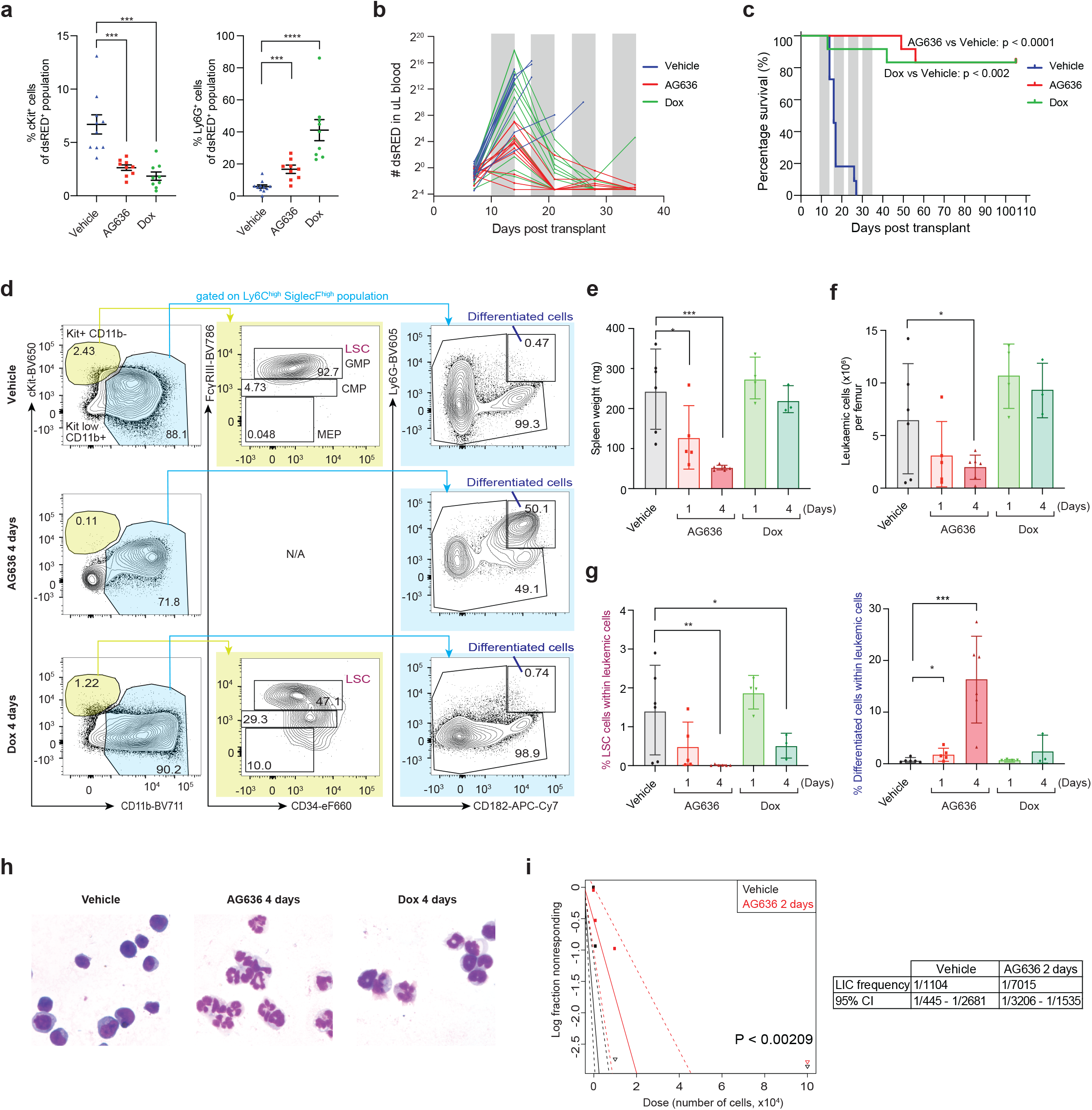
AG636 is an effective single agent therapy in MLL-rearranged AML. **a**, Frequency of MN leukemic cells expressing the immature marker cKit and mature myeloid marker Ly6G in the peripheral blood following 5 days of treatment (*n =* 9-10 mice/group; mice with <2% tumor burden in any condition were censored from the analysis). **b**, Absolute number of MN cells in the peripheral blood quantified by flow cytometry. Grey bars denote treatment (*n =* 11-12 mice/group). **c**, Kaplan-Meier survival curve of leukemic mice (*n* = 11-12 mice/group, median survival is 16.5 for vehicle and not reached for AG636 and doxycycline, p < 0.0001 by Log-rank test). **d**, Representative FACS plots showing differentiation induced by AG636 and doxycycline. **e**, Spleen weights of MN tumor-bearing mice. **f**, absolute number of MN cells in the bone marrow. **g**, Frequency (as percentage of all MN cells) of LSCs (CD11b^low^cKit^high^FcgR^+^) and differentiated cells (CD182^+^Ly6G^+^) in the bone marrow (*n =* 3-6 mice/group). **h**, May-Grunwald-Giemsa-stained cytospins of sorted MN cells showing myeloid differentiation. **i**, Quantification of functionally-define LSCs using a limiting dilution assay calculated by ELDA (*n* = 4-8 mice/group). Data in **a** and **e-g** are presented as mean ± s.d.; *P* values were calculated using a one-tailed Student’s unpaired t-test. \**P* < 0.05, \*\**P* < 0.01, \*\*\**P* < 0.001; Dox - doxycycline.

To characterize the cellular response of MN cells that underpins the therapeutic efficacy of DHODH inhibition, we next performed additional short-term treatments. Mice bearing MN tumors were treated with doxycycline or AG636 for one or four days and detailed immunophenotyping was performed on the peripheral blood, spleen and bone marrow compartments (Fig. 1d-g and Extended Data Fig. 1e-f). As expected, doxycycline rapidly silenced MLL-AF9 expression (not shown) and induced myeloid differentiation of MN cells, evidenced by a reduction in the frequency and absolute number of phenotypically-defined LSCs (CD11b^low^cKit^high^FcγR3^+^)^16^, and a concomitant increase in differentiated granulocyte-like cells (CD182^+^Ly6G^+^)(Fig. 1g and Extended Data Fig. 1f). DHODH inhibition similarly induced myeloid differentiation, but notably the phenotype was different to that induced by genetic depletion of MLL-AF9. In particular, AG636 rapidly reduced splenomegaly as well as the total number of leukemic cells in the peripheral blood, spleen and bone marrow of drug-treated animals (Fig. 1e-g and Extended Data Fig. 1e-f). Consistent with the flow cytometric data, morphological analysis of sorted MN cells revealed the presence of myeloid maturation in both doxycycline and AG636-treated mice (Fig. 1h). Importantly, the viable MN cells remaining after two days of AG636 therapy possessed reduced leukemia initiating capacity in limiting dilution re-transplant experiments, demonstrating functional loss of LSCs (Fig. 1i). Overall, AG636 has excellent potency against MLL-rearranged AML, effectively targets LSCs and induces rapid disease regression through a combination of cell death and differentiation.

### Inhibition of *de novo* pyrimidine synthesis is broadly effective against multiple AML subtypes

To determine whether AG636 has efficacy in other subtypes of AML we tested two additional murine models that carry distinct combinations of AML driver genes using an endpoint of tumor burden following four days of treatment. Recipient mice were engrafted with established tumors expressing the core binding factor fusion protein RUNX1-RUNX1T1 (also known as AML1-ETO) ^17^ or the combination of mutant alleles of *IDH1*^*R132H*^, *DNMT3A*^*R882H*^ and *Nras*^*G12D*^ (hereafter referred to as I1DN, see Extended Data Fig. 2a and Methods for details) and randomized to receive vehicle or AG636 therapy. Both models co-expressed fluorescent reporters, enabling precise compartment-specific quantification of leukemic burden. In both contexts, we observed reduced tumor progression in the drug group (Fig. 2a-c). In the RUNX1-RUNX1T1 model, the spleen weights of AG636-treated mice were significantly reduced and although the overall number of leukemic cells in the bone marrow were comparable to controls, fewer cells expressed cKit and more cells expressed CD11b demonstrating differentiation (Fig. 2a-d and Extended Data 2b-c). In the I1DN model, AG636 reduced the spleen weight as well as the total number of leukemic cells in both the spleen and bone marrow (Fig. 2a-c). Unlike in the MN and RUNX1-RUNX1T1 AMLs, there was minimal impact on expression of cKit and CD11b, suggesting that the predominant effect of AG636 in the I1DN setting was to inhibit proliferation and/or trigger cell death (Fig. 2e and Extended Data Fig. 2d-e). Interestingly, in both MN and I1DN models we observed increased surface expression of Sca1, a marker associated with stemness but also with Type I interferon (IFN) signaling ^18^(Extended Data Fig. 2c and 2e). Taken together, our data demonstrate that the therapeutic potential of DHODH inhibition in AML is not limited to individual disease subsets.

**Fig. 2:**
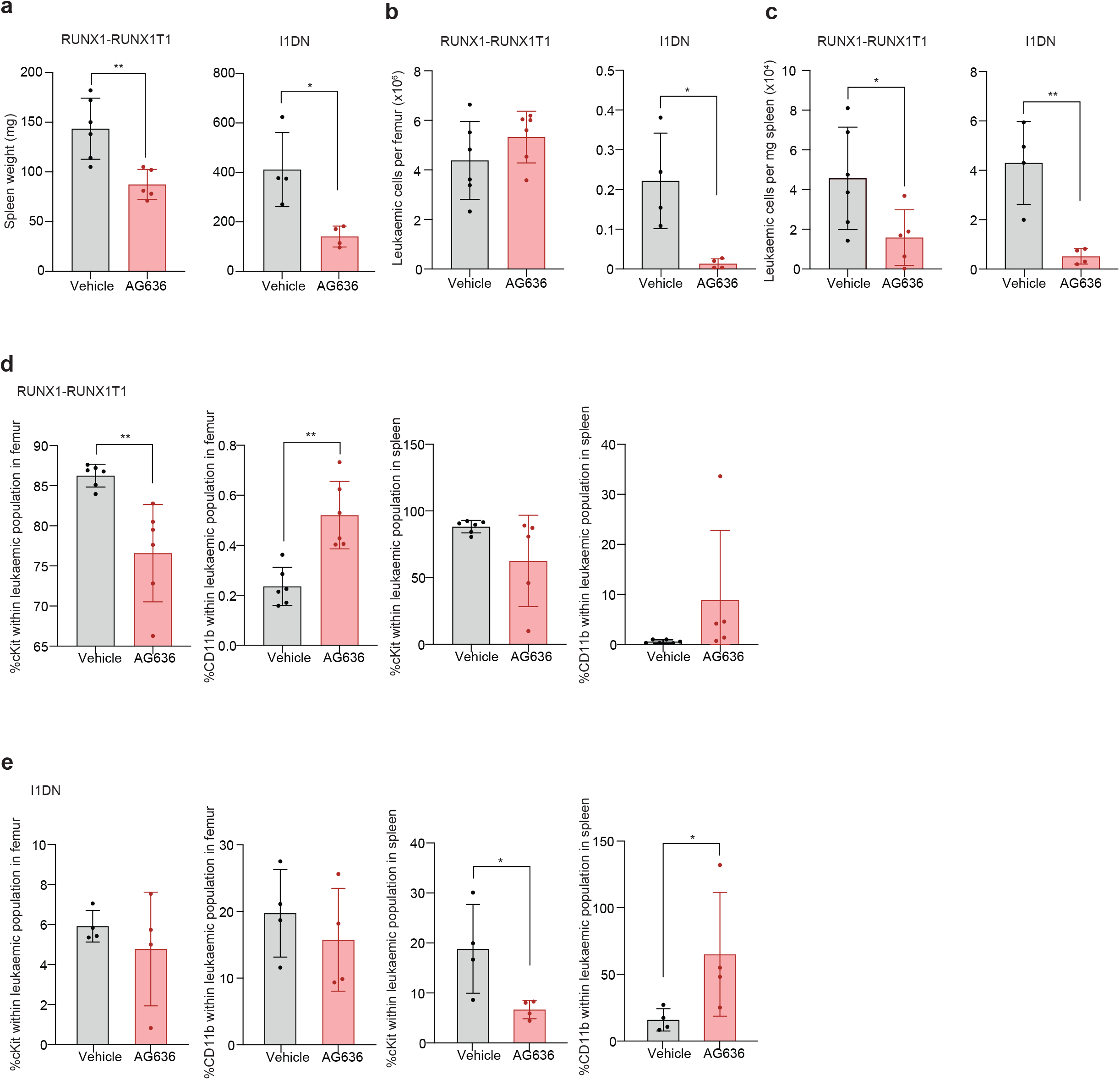
DHODH inhibition induces differentiation and inhibits proliferation in multiple AML sub-types. **a**, Spleen weights of RUNX1-RUNX1T1 or I1DN tumor bearing mice treated with AG636 or vehicle for 4 days. **b-c**, Absolute number of leukemic cells in the bone marrow (**b**) and spleens (**c**) of RUNX1-RUNX1T1 or I1DN tumor bearing mice treated with AG636 or vehicle for 4 days. **d-e**, Frequency of leukemic cells expressing the immature marker cKit and mature myeloid marker CD11b in the bone marrow and spleens of RUNX1-RUNX1T1 (**d**) or I1DN (**e)** tumor bearing mice treated with AG636 or vehicle for 4 days (*n =* 6 mice/group). *n =* 5-6 mice/group for RUNX1-RUNX1T1 model, *n* = 4 mice/group for I1DN model; data are presented as mean ± s.d.; *P* values were calculated using a one-tailed Student’s unpaired t-test. \**P* < 0.05, \*\**P* < 0.01, \*\*\**P* < 0.001.

### Effects of DHODH inhibition on normal hematopoiesis

The clinical utility of potential anti-cancer agents that target proteins which are expressed in non-malignant cells is dependent on the existence of a therapeutic window. As DHODH is widely expressed in healthy hematopoietic cells, we sought to assess the impact of AG636 on normal blood development *in vivo*. We treated non-tumor bearing mice with the same regimen that had a significant impact on leukemia development and quantified stem, progenitor and mature cell populations from all three major hematopoietic lineages (lymphoid, myeloid, erythro-megakaryocytic)(Fig. 3a). Overall the consequences of DHODH inhibition on normal blood and bone marrow populations were minor in comparison with the effects observed on AML cells (Fig. 3, Extended Data Fig. 3 and Extended Data Table 1). Drug treatment did not affect peripheral red or white blood cell counts (Fig. 3b-c), but resulted in a minor reduction in platelet counts (Fig. 3d) and a reduced overall number of bone marrow mononuclear cells (MNCs)(Fig. 3e). The latter was attributable to an approximately 50% reduction in B cells, whereas myeloid, T and NK cells were largely unaffected (Fig. 3f). Within the erythroid compartment, the immature ProE, EryA, and EryB fractions were reduced by drug treatment, whereas the EryC fraction was increased, suggesting a burst of erythroid differentiation (Fig. 3g). Within the hematopoietic stem/progenitor (HSPC) compartment, the greatest impact was on myeloid progenitors, with a reduction in common myeloid progenitors and megakaryocyte-erythroid progenitors (Fig. 3h-i). The overall number of multi-potent progenitors and short-term hematopoietic stem cells was not affected, although there was a reduction in long-term hematopoietic stem cells (Fig. 3j-k). Collectively, these data are consistent with the observation that as many as four weekly cycles of AG636 treatment are well tolerated even in sub-lethally irradiated tumor-bearing mice and point to a selective vulnerability of AML cells to pyrimidine starvation.

**Fig. 3:**
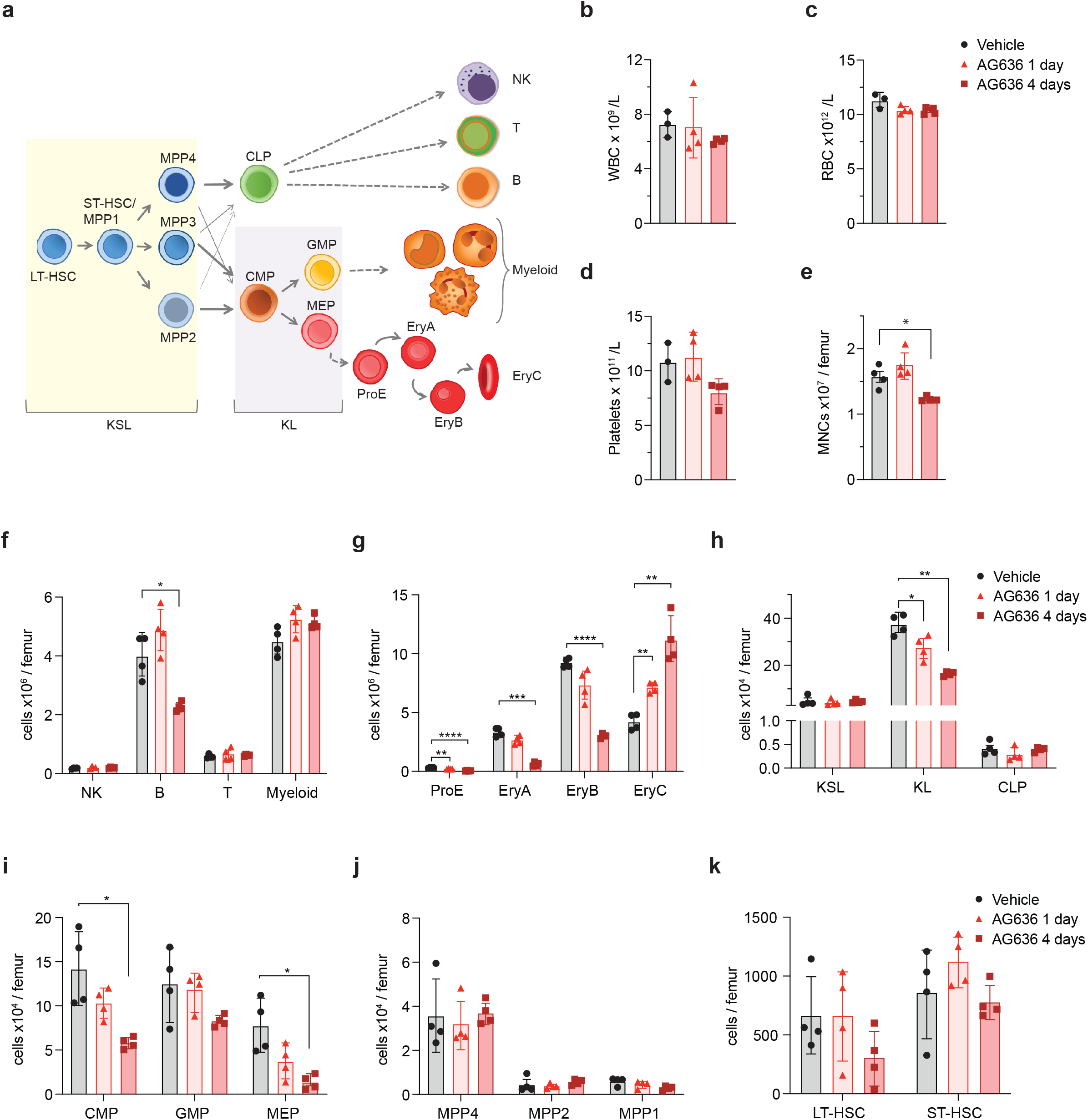
DHODH inhibition has minor impact on normal blood development. **a**, Schematic of hematopoietic differentiation. **b-d**, Peripheral blood white blood cell (WBC, **b**), red blood cell (RBC, **c**) and platelet counts (**d**) quantified by Sysmex cell counter in mice treated with AG636 or vehicle for 1 or 4 days (*n =* 4 mice/group). **e-k**, Various bone marrow populations were quantified by flow cytometry in mice treated with AG636 or vehicle for 1 or 4 days (see Extended Data Table 1 for abbreviations and markers). *n =* 4 mice/group; data are presented as mean ± s.d.; *P* values were calculated using a two-tailed Student’s unpaired t-test; only comparisons that meet the threshold of *P* < 0.05 are shown. \**P* < 0.05, \*\**P* < 0.01, \*\*\**P* < 0.001.

### AG636 downregulates protein synthesis pathways in LSCs

LSCs play a central role in AML pathogenesis and their metabolic requirements and gene expression programs are distinct from those of “bulk” leukemic cells ^12,19^. To gain insights into the molecular pathways that are perturbed by DHODHi in LSCs we performed RNA sequencing (RNAseq). We initially focused on the MN model, and compared the effects of AG636 to differentiation induced by genetic depletion of MLL-AF9. To that end, we sorted and analyzed cKit^high^CD11b^low^ immature MN cells that are highly enriched in functional LSCs^16^ from mice treated with vehicle, AG636 for one day, or doxycycline for one or four days. We were unable to perform analysis following four days of AG636 treatment due to insufficient numbers of cKit^high^CD11b^low^ MN cells.

Using stringent statistical cut-offs, we identified differentially expressed genes (DEGs) in all treatment conditions (Extended Data Table 2). As expected, doxycycline induced robust transcriptional changes, particularly at the four-day time-point. Consistent with previous studies, we observed perturbation of classic MLL-AF9-regulated genes including *Myb, Hoxa5, Hoxa9, Tcf4 and Id2* ^14,15^. Altered gene expression induced by AG636 partially overlapped with changes induced by MLL-AF9 depletion, although 47% of DEGs (99/231 genes up-regulated by AG636 and 139/275 genes down-regulated by AG636) were unique to DHODH inhibitor treatment (Fig. 4a). We performed gene set enrichment analysis ^20^ to identify pathways that may be common or unique between the treatment conditions. Confirming our phenotypic observations, gene signatures associated with myeloid differentiation were highly positively enriched in both treatment conditions. In contrast, gene sets related to protein translation (down-regulated) and type I interferon signaling (up-regulated) were uniquely enriched in AG636-treated cells (Fig. 4b). Of note, inhibition of *de novo* pyrimidine synthesis has previously been reported to trigger expression of interferon stimulated genes^21^ and type I interferons have been implicated in differentiation of LSCs^22^.

**Fig. 4:**
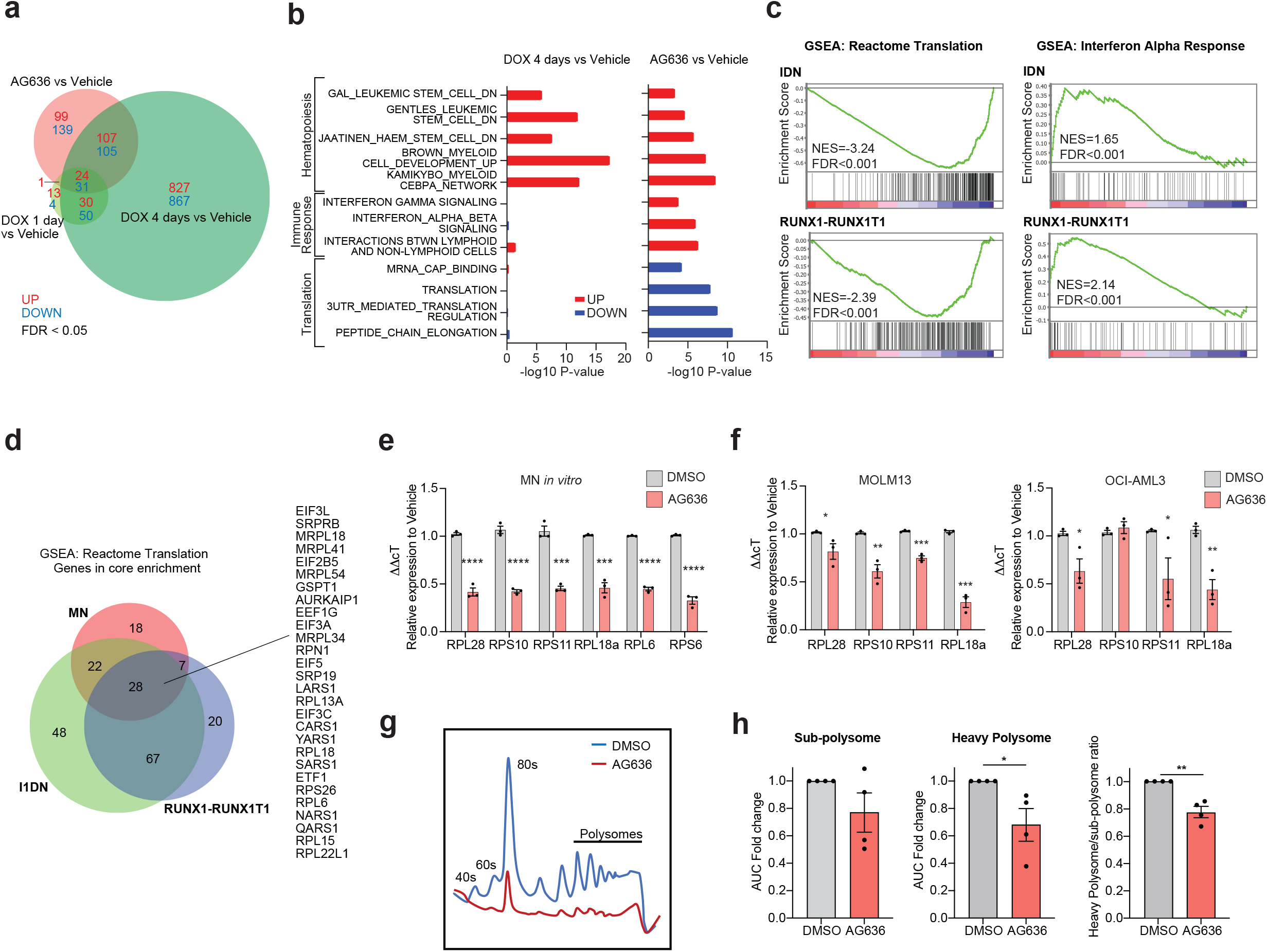
AG636 induces transcriptional downregulation of genes required for protein translation. **a-b**, RNA sequencing performed on cKit^high^CD11b^low^ MN cells sorted from AG636, doxycycline or vehicle treated mice (*n =* 3 mice/group). **a**, Venn diagram showing overlap in DEGs between different treatment conditions. **b**, Gene set enrichment analysis showing common and differential enrichment of biological pathways in gene expression data from AG636 or doxycycline treated animals. Gene sets are from C2:CGP and Reactome sub-collections in MSigDB database (see methods for more information). **c**, RNA sequencing performed on cKit^+^CD11b^-^ RUNX1-RUNX1T1 or I1DN cells sorted from mice treated with AG636 or vehicle for 1 day (*n =* 3 mice/group). Bar code plots showing enrichment of selected pathways. **d**, Venn diagram showing overlap of genes in the core enrichment within the Reactome Translation gene set in the MN, RUNX1-RUNX1T1 and I1DN models. **e**,**f** qPCR showing down-regulation of genes encoding ribosomal proteins in MN cells (**e**) and human AML cell lines (**f**) treated for 24 hours with AG636 (*n =* 3 biological replicates for each cell line). **g-h**, Polysome profiling of MN cells treated *in vitro* with AG636 or vehicle for 24h. Representative trace (**g**) and quantification of sub-polysome and heavy polysome fractions (**h**) determined by measuring area under the curve (AUC) (*n =* 4 biological replicates). Data in **e, f** and **h** are presented as mean ± s.e.m.; *P* values were calculated using a one-tailed Student’s unpaired t-test. \**P* < 0.05, \*\**P* < 0.01, \*\*\**P* < 0.001, ****P<0.0001.

Recent studies have demonstrated that LSCs from different patients and model systems share many common properties despite their unique collection of driver mutations ^23–26^. To determine whether the transcriptional effects of AG636 were conserved in non-MN LSCs, we analyzed LSC-enriched populations from the RUNX1-RUNX1T1 and I1DN models (cKit^high^ and cKit^+^CD11b^-^ cells, respectively). We found that DHODHi perturbed similar pathways in all three contexts (Fig. 4c and Extended Data Tables 3 and 4), although we also detected some model-specific changes (e.g. transforming growth factor β (TGF-β) signaling and RNA polymerase I components)(Extended Data Fig. 4a-b). Overall, the magnitude of change was greatest for transcripts encoding ribosomal proteins and translation initiation factors such as *Rpl15, Eif5* and *Eif3A* (Fig. 4d). Interestingly, mRNAs that encode these proteins have been shown to have a relatively long half-life in mammalian cells ^27,28^. In contrast, transcripts with a short half-life such as *Myc* were not affected by AG636 treatment. Moreover, unbiased analysis using a dataset of mRNA stability from murine NIH3T3 fibroblasts ^28^ revealed no consistent trends in perturbation of short-lived transcripts (Extended Data Fig. 4d). Consistent with our *in vivo* data in LSCs, we found that ribosomal protein genes were downregulated in MN cells and human AML cell lines treated *in vitro* with AG636 (Fig. 4e-f). Importantly these changes occurred within 24 hours of drug administration and preceded loss of viability of the cells (Extended Data Fig. 4d).

Finally, to determine whether the transcriptional perturbations affected the assembly of ribosomal subunits we performed polysome profiling. As these experiments require a large amount of material, we treated MN cells *in vitro* with AG636. As expected, DHODH inhibition caused a profound reduction in heavy polysomes as well as the heavy polysome to sub-polysome ratio (Fig. 4g-h). Altogether, these findings suggest that down-regulation of protein synthesis pathways is a specific and early response of AML cells to DHODH inhibition that occurs at the level of mRNA synthesis and is not attributable to global dampening of transcription caused by reduced availability of pyrimidine nucleotides.

### Analysis of chromatin accessibility identifies YY1 as a key transcription factor that modulates gene expression downstream of DHODH inhibition

To identify transcription factors (TFs) that drive altered gene expression in AML cells in response to pyrimidine starvation *in vivo* we mapped open chromatin using the assay for transposase-accessible chromatin (ATACseq). We concentrated our analysis on the MN model and specifically on the cKit^high^CD11b^low^ population that is highly enriched in LSCs. We identified approximately 150,000 peaks that were distributed at or near transcription start sites (TSS), within gene bodies or intergenic regions (Fig. 5a and Extended Data Fig. 5a). As expected, TSS-associated open regions were strongly correlated with gene expression (Extended Data Fig. 5b). AG636 induced selective alterations to chromatin accessibility, with approximately 2% of all ATAC peaks showing differential signal between drug and vehicle treated cells (Fig. 5a). Most of the differential peaks were present in intergenic regions, suggesting that AG636 treatment alters enhancer utilization (Fig. 5b). Concordant with our morphological and gene expression findings, regions of increased accessibility induced by DHODH inhibition were enriched in DNA motifs for known myeloid differentiation factors including PU.1 and members of the CEBP family (Fig. 5c). These regions were also enriched for ATF4 motifs, suggesting that drug treatment activated the integrated stress response in LSCs as previously reported in colorectal cancer cells following mitochondrial complex III inhibition^29^.

**Fig. 5.**
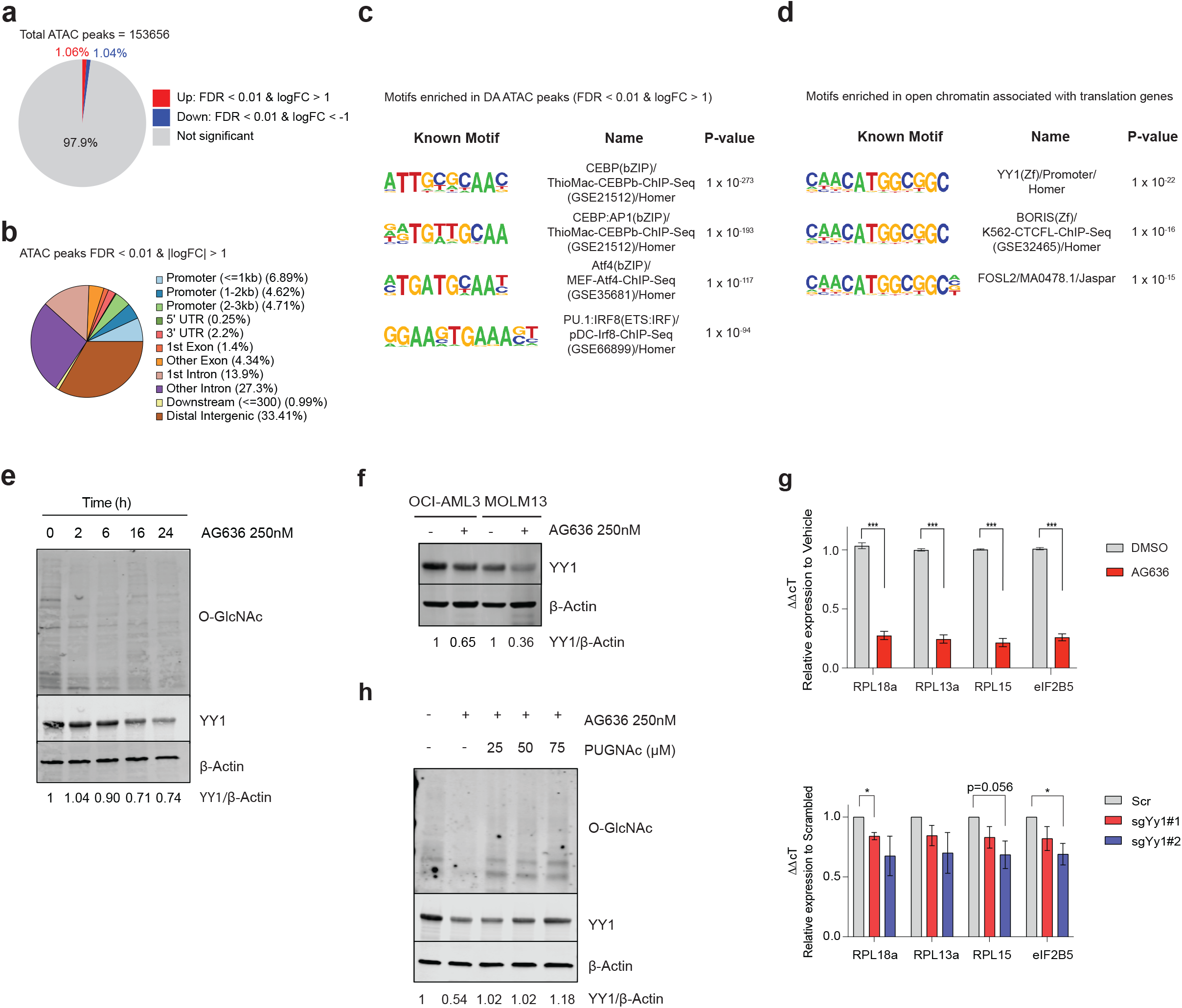
Downregulation of YY1 protein contributes to altered gene expression in AML cells following DHODH inhibition. **a-d**, ATAC sequencing performed on cKit^high^CD11b^low^ MN cells sorted from mice treated with AG636 or vehicle for 2 days (*n =* 3 mice/group). **a**, Pie chart showing the relative proportion of ATAC peaks where chromatin accessibility is increased, decreased or unchanged by DHODH inhibition. **b**, Pie chart showing association between regions of differential chromatin accessibility and different genome features. **c**, HOMER motif analysis showing transcription factor motifs enriched within regions of differential chromatin accessibility. **d**, HOMER motif analysis showing enrichment of YY1 motifs within regions of accessible chromatin associated with translation genes. All other regions of open chromatin were used as the background. **e**, Time-course showing global down-regulation of O-GlcNAcylation and YY1 expression in MN cells treated with AG636. The experiment was repeated twice with similar results. **f**, Western blot of YY1 expression in human AML cell lines treated with AG636 for 24 hours **g**, qPCR showing down-regulation of genes encoding ribosomal proteins that are putative YY1 targets in Cas9 expressing MN cells treated with AG636 or DMSO (top); or transduced with sgRNAs targeting YY1 or control sgRNAs (bottom) (*n =* 2 biological replicates). Data are presented as mean ± s.e.m.; P values were calculated using one-tailed Student’s unpaired t-test. *P < 0.05,\*\*\**P* < 0.001. **h**, Western blot in MN cells co-treated with AG636 and PUGNAc for 24 hours.

We next focused our attention on genes involved in protein translation (hereafter “translation genes”), as the coordinated down-regulation of this pathway represents the most striking and conserved phenotype triggered by AG636 that we observed in AML LSCs *in vivo*. We used the Pscan algorithm to identify over-represented TF binding sites within the proximal promoters of translation genes^30^. This analysis identified members of the YY and ETS families as the most highly enriched (Extended Data Table 5). YY motifs were also highly enriched in translation gene-associated regions of open chromatin compared with other regions of open chromatin in MN LSCs (Fig. 5d). YY1, but not YY2 was highly expressed in all three murine AML models, as well as in publicly available human AML gene expression data (Extended Data Fig. 5c) ^31^. Notably, YY1 has previously been implicated as a transcriptional activator of the translation pathway ^32^ and its activity is known to be regulated by O-Linked N-Acetylglucosaminylation (O-GlcNAc), a common protein post-translational modification that requires pyrimidine synthesis ^7,33,34^.

Confirming previous reports^7,9^, we found that DHODH inhibition rapidly reduced global O-GlcNAc protein modification in MN cells (Fig. 5e). Loss of O-GlcNAc coincided with down-regulation of YY1 protein (Fig. 5e). AG636 treatment similarly caused downregulation of YY1 protein, but not transcript, in human AML cell lines MOLM13 and OCI-AML3 (Fig. 5f and Extended Data Fig. 5d). Notably, the phenotype was more pronounced in MOLM13 cells, which are more sensitive to AG636-induced killing compared with OCI-AML3 cells where the effects of the inhibitor at early time points are predominantly cytostatic (Extended Data Fig. 4d).

We next used CRISPR to disrupt *YY1* in MN cells using two independent sgRNAs. YY1 is a common essential gene as defined by the Cancer Dependency Map (DepMap) ^35^ and MN cells containing *YY1*-targeting sgRNAs and co-expressing a Crimson (Crim) fluorescent reporter were depleted over time in culture (Extended Data Fig. 5e). To determine the impact of YY1 deletion on expression of translation genes we sorted viable newly transduced cells on day three post viral infection by which time we observed substantial reduction of YY1 protein expression (Extended Data Fig. 5f). qPCR analysis focused on a subset of translation genes that contained a YY1 motif within their promoter confirmed that YY1 knockout caused downregulation of these transcripts as expected (Fig. 5g).

O-GlcNAc modification is regulated by the reciprocal activity of two evolutionary conserved enzymes - O-GlcNAc transferase (OGT) which deposits the mark, and O-GlcNAcase (OGA) which removes it^34^. We tested whether inhibition of OGA could rescue AG636-mediated YY1 protein downregulation. To that end, we co-treated MN cells with AG636 and PUGNAc, an inhibitor of OGA. Concordant with our hypothesis, PUGNAc countered the loss of YY1 protein caused by AG636 (Fig. 5h). Thus, our data suggest that downregulation of YY1 protein downstream of DHODH inhibition underpins the downregulation of the protein translation pathway following drug treatment.

### Loss of CDK5 and INO80 complex sensitizes AML cells to inhibition of *de novo* pyrimidine synthesis

Pooled CRISPR screens are a powerful approach to systematically determine key factors that mediate drug resistance and sensitivity. To complement our analyses of the molecular and cellular responses of AML cells to DHODHi we performed a CRISPR knockout screen to uncover genes that increase or decrease treatment efficacy. As epigenetic regulators are frequently dysregulated in AML^31^ and many are amenable to therapeutic targeting with existing molecules^36^ we focused specifically on this class of genes. MN cells were transduced with an epigenetics-targeted sgRNA library, cultured in DMSO or increasing concentrations of AG636 and the relative distribution of sgRNAs at the beginning of the experiment and after 10 and 24 days of culture quantified by sequencing (Fig. 6a). We ranked positively and negatively selected sgRNAs and genes using the MAGeCK algorithm^37^ and integrated data across different conditions and timepoints.

**Fig. 6.**
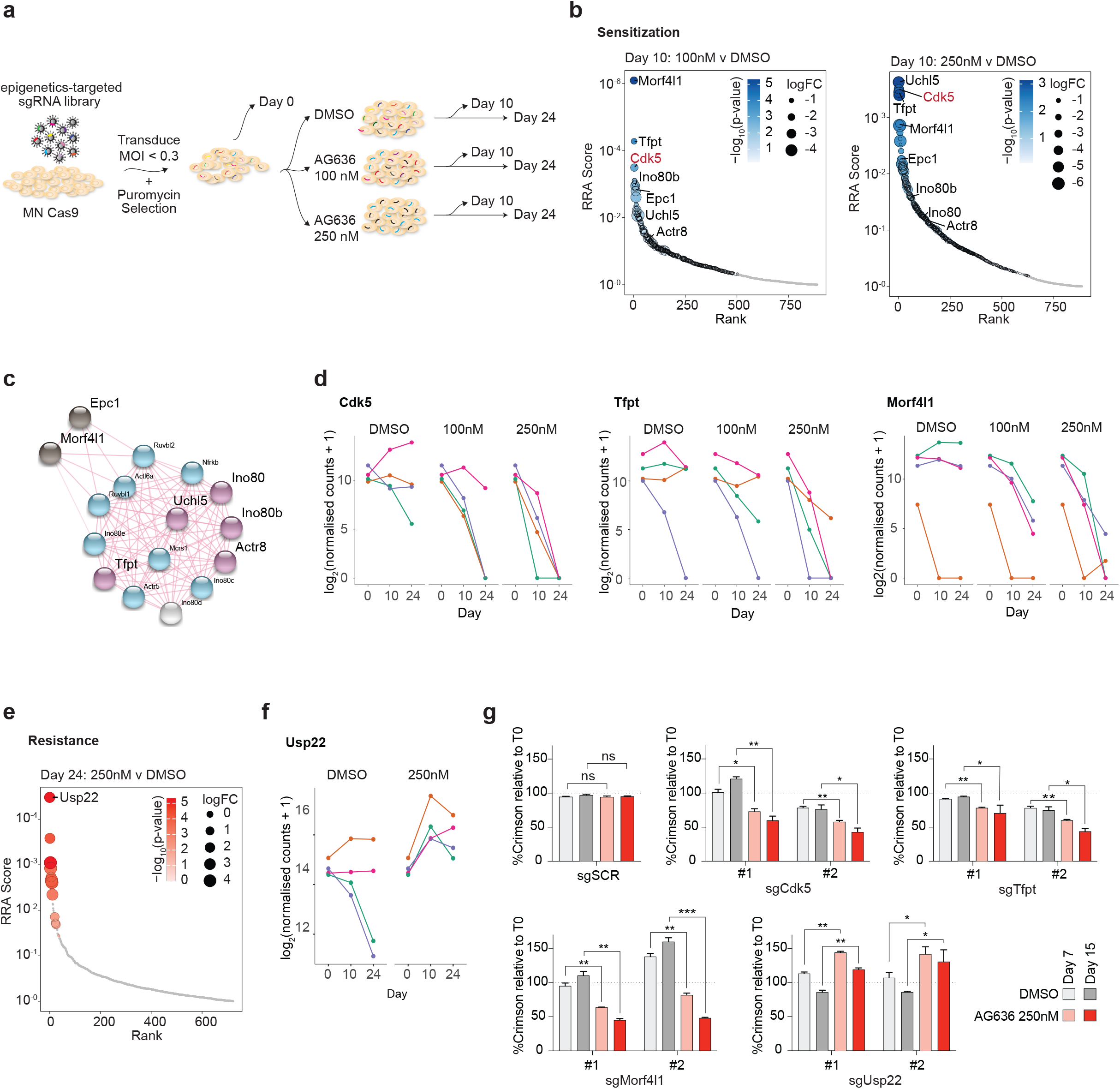
Pooled CRISPR screen identifies CDK5 and INO80 complex as modulators of AG636 sensitivity. **a**, Schematic of the pooled CRISPR screen in MN AML cells. **b**, Rank plot of MAGeCK analysis showing genes that were negatively selected at day 10 in cells treated with 100nM or 250nM AG636. **c**, STRING network analysis showing known interactions between components of the INO80 complex and associated proteins. Blue - INO80 complex genes which are essential (negatively enriched in all conditions); light grey - INO80 complex gene which was positively enriched in all conditions; purple - INO80 complex genes which are AG636 sensitizers (negatively enriched in AG636 condition only); black - other AG636 sensitizers identified in the screen. **d**, Normalized counts for sgRNAs targeting Cdk5, Tfpt and Morf4l1. **e**, Rank plot of MAGeCK analysis showing genes that were positively selected at day 24 in cells treated with 250nM AG-636. **f**, Normalized counts for sgRNAs targeting Usp22. **g**, Proliferative competition assays in MN cells transduced with individual sgRNAs and cultured in AG636 or DMSO (*n =* 2 biological replicates; data represented as mean ± s.e.m; P values were calculated using one-tailed Student’s unpaired t-test. \**P* < 0.05, \*\**P* < 0.01, \*\*\**P* < 0.001).

To identify genes that were negatively selected in the presence of drug we concentrated on day 10 at which time the relative abundance of most sgRNAs compared with day 0 was unaffected (Extended Data Fig. 6a). One of the most prominent genes in this analysis was Cdk5 with all 4 sgRNAs in the library being lost in a drug concentration and time-dependent manner (Fig. 6b and d). Additionally, reinforcing our findings implicating YY1 as a major effector of the transcriptional response to DHODH inhibition, many of the other top ranked sensitizer hits were components of the INO80 chromatin remodeling complex. This included core proteins Tfpt, Actr8, Ino80b and Uchl5 as well as associated proteins Morfl1 and Epc1 (Fig. 6b-d and Extended Data Fig. 6b). INO80 is physically associated with YY1 and is recruited to YY1 target loci to activate gene expression^38^. The INO80 factors that did not synergize with AG636 treatment were essential for the proliferation of MN cells and analysis of DepMap data ^35^ confirmed that these genes were broadly essential in AML (Extended Data Fig. 6b-c).

Conversely, positive selection was most obvious at day 24 in the high AG636 concentration when many sgRNAs were depleted (Extended Data Fig. 6a). The histone deubiquitinase Usp22 was the most significantly positively enriched gene in our library with the representation of all 4 guides being significantly increased in drug treated cells compared with both day 0 and DMSO conditions (Fig. 6e-f).

We further validated the phenotype of a subset of genes identified in the screen using two sgRNAs per gene in individual competition assays. As expected, Usp22 targeting sgRNAs mediated resistance to AG636, whereas Cdk5, Tfpt and Morfl1 guides synergized with drug treatment (Fig. 6g). Thus, we have uncovered novel factors that have not previously been implicated in regulating the sensitivity of AML cells to DHODH inhibition.

### Inhibition of CDK5 and DHODH has synergistic activity in AML

One of the top sensitization hits that was revealed in our screen was CDK5. In the human genome CDK5 is localized on the long arm of chromosome 7 that is recurrently deleted in AML and myelodysplastic syndrome (MDS) and associated with poor prognosis ^39^. Numerous clinical-grade molecules that inhibit CDK5 have also been developed. These two factors provided a strong impetus to further explore the interaction between CDK5 loss and DHODH inhibition. Consistent with our observations in MN cells, CDK5-targeting guides also synergized with AG636 treatment in human AML cell lines MOLM13 and OCI-AML3 (Fig. 7a). Notably, CDK5 was dispensable for cellular proliferation and did not mediate sensitization to the DNA damaging agent cytarabine (AraC), demonstrating the selective nature of the phenotype (Fig. 7a and Extended Data Fig. 7a).

**Fig. 7.**
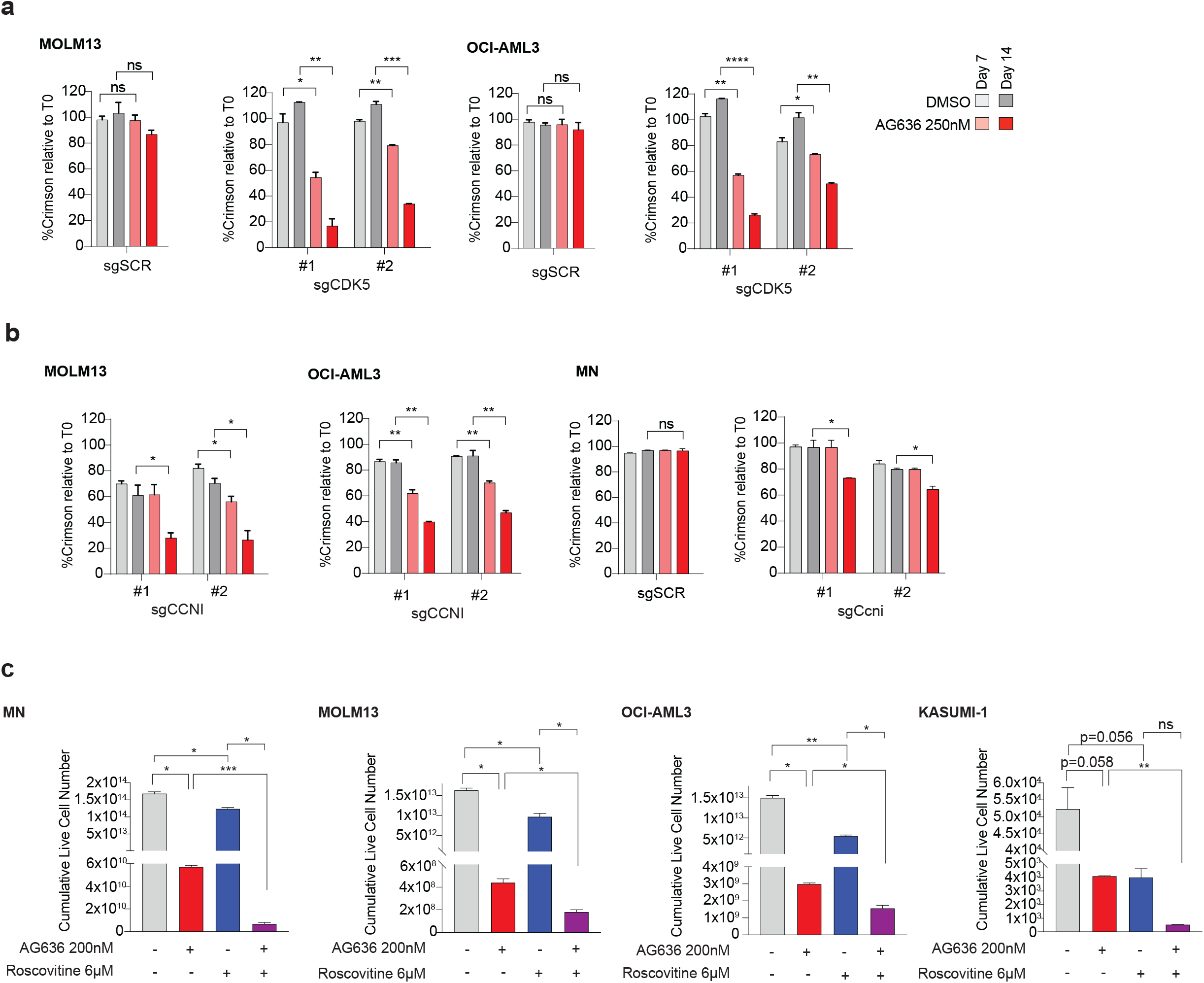
Genetic or pharmacological inhibition of CDK5 increases sensitivity of AML cells to blockade of pyrimidine synthesis. **a-b**, Proliferative competition assays in human AML cell lines transduced with indicated sgRNAs and cultured in AG636 or DMSO (*n =* 2 biological replicates; data represented as mean ± s.e.m; P values were calculated using one-tailed Student’s unpaired t-test. \**P* < 0.05, \*\**P* < 0.01, \*\*\**P* < 0.001, ***P<0.0001). **c**, AML cells were treated with AG636, Roscovitine or the combination for 17 days. Cumulative live cell counts are shown (*n =* 2 biological replicates; data represented as mean ± s.e.m; P values were calculated using Brown-Forsythe and Welch ANOVA test. \**P* < 0.05, \*\**P* < 0.01, \*\*\**P* < 0.001).

CDK5 is an atypical member of the cyclin-dependent kinase family the catalytic activity of which can be activated by Cyclin I (*CCNI*), but also non-cyclin co-activators p35 (*CDK5R1*) and p39 (*CDK5R2*) ^40,41^. We analyzed gene expression data from AML patients in The Cancer Genome Atlas dataset^31^and found that *CDK5, CCNI* and *CDK5R1* are robustly expressed, whereas *CDK5R2* mRNA could not be detected in the majority of samples (Extended Data Fig. 7b). In competition assays, *CCNI*-targeting sgRNAs similarly sensitized AML cells to AG636 treatment, whereas *CDK5R1* knockout had no effect (Fig. 7b and Extended Fig. 7c). Thus, our data implicates CDK5/CCNI as synthetic lethal partners with DHODH in AML.

We then tested whether AG636 could synergize with CDK5 inhibitors. Most existing CDK5 inhibitors show poor selectivity for CDK5 over other cyclin-dependent kinases due to their high sequence homology. We used Roscovitine, an ATP-competitive inhibitor that has preferential activity against CDK5, but also blocks CDK1, 2, 7 and 9 ^42,43^, and performed drug synergy assays in AML cell lines. Co-treatment with AG636 and Roscovitine greatly increased cell death, especially in MLL-rearranged MN and MOLM13 cells where Roscovitine alone had only mild effects (Fig. 7c). Given that CDK5 expression is dispensable in these cells, whereas CDK1, 2, 7 and 9 are common essential genes required for the proliferation of almost all cancer cells lines (Extended Data Fig. 7d), the effects of Roscovitine are highly likely to be attributable to inhibition of CDK5. These results provide further evidence that CDK5 is a biomarker for the efficacy of DHODH inhibition in AML and provide proof of principle for a novel combinatorial strategy.

## Discussion

Many tumor cells reprogram their metabolism to meet altered demands for macromolecules and energy. Small molecules that interfere with DNA or RNA synthesis have been extensively used in cancer chemotherapy, demonstrating that although these processes are ubiquitous, they can nonetheless be targeted for therapeutic benefit. Critically however, improvements to patient outcomes require a more sophisticated strategy. To that end, studies over the past decade have sought to identify selective metabolic vulnerabilities of cancer cells. Notably, an altered metabolic state appears to be a common feature of AML LSCs that is broadly conserved across different genetic contexts and is emerging as a potential Achilles’ heel in leukemia ^23–26^.

The discovery linking pyrimidine starvation to cell fate in AML has reinvigorated interest in DHODH as an anti-cancer target and has spurred extensive efforts to develop more potent and selective next-generation inhibitors ^7–11^. In agreement with previous studies, we found that DHODH inhibition has excellent potency in different AML sub-types *in vivo*. The results were particularly striking in the highly aggressive MLL-rearranged MN model, where most conventional or targeted agents induce only partial short-lived responses ^13,44,45^. In contrast, AG636 induced long-lasting remission in almost all treated mice and these responses were sustained even after drug withdrawal. Drug treatment caused a mix of cell death and differentiation of leukemic cells leading to a rapid reduction in tumor burden and loss of LSC activity as confirmed by both morphological analysis and functional assays. The relative proportion of cell death and differentiation varied between the different AML sub-types and even between different compartments within individual animals, suggesting that phenotypic differentiation may not be an effective biomarker of drug response in all contexts.

The use of syngeneic tumor models enabled us to directly compare the sensitivity of malignant and non-malignant hematopoietic cells to DHODHi *in vivo*. Although, DHODH is widely expressed in healthy tissues, the effects of AG636 on non-malignant blood and bone marrow cells were minor in comparison with the effects observed on AML cells. Interestingly, of all the mature, progenitor and stem cell populations analyzed, drug treatment had the greatest impact on B-cells, reducing their total number in the bone marrow by ∼50%. These findings correlate with the apparent exquisite dependence of malignant B-cells on DHODH expression and suggest that the B-cell lineage may be intrinsically sensitive to DHODHi ^11^. Importantly, B-cell aplasia is tolerable within the context of cancer therapy and can be treated by immunoglobulin replacement.

Knowledge of prognostic biomarkers that predict or accompany an effective therapeutic response is essential for patient selection and effective clinical deployment of anti-cancer drugs. This is especially pertinent for drug targets such as DHODH that are ubiquitously expressed and rarely mutated or over-expressed in cancer. We found that transcriptional downregulation of translation genes is a common event following DHODHi treatment that occurs independently of differentiation and is conserved across multiple AML sub-types. Mechanistically, our data suggest that the transcription factor YY1 and the associated INO80 chromatin remodeling complex play critical roles in this molecular response. The response is triggered at least partly because inhibition of pyrimidine synthesis downregulates protein O-GlcNAcylation leading to loss of YY1 protein. O-GlcNAc is a common modification that regulates the stability, trafficking and function of many proteins and OGT, the sole enzyme known to catalyze O-GlcNAcylation, is indispensable for the proliferation of all 975 cancer cell lines in the DepMap database ^34,35^. Hence, it is likely that global loss of GlcNAc downstream of DHODHi exerts multiple anti-proliferative effects.

Our targeted CRISPR screen also revealed a hitherto unknown relationship between inhibition of pyrimidine synthesis and CDK5. CDK5 has been most extensively characterized in the nervous system and has been implicated in the pathogenesis of Alzheimer’s and Parkinson’s diseases^40^. More recently, numerous studies have described a role for CDK5 in non-neuronal cells including in AML where it was shown to regulate the pro-apoptotic protein NOXA following glucose deprivation^41^. Our study demonstrates that genetic or pharmacological ablation of CDK5 increases DHODHi efficacy in AML. As loss of chromosome 7 (−7/7q) that encodes CDK5 is a poor prognostic marker in myeloid malignancies, our results open the door to prioritize clinical evaluation of DHODHi in a challenging patient population for which there are currently limited therapeutic options ^39^.

In conclusion, our study provides novel insights into the activity of DHODH inhibitors in AML and adds to the growing body of evidence supporting further development of these compounds. Importantly, the response biomarkers that we have identified can aid these efforts and ensure that clinical trials are appropriately powered.

## Methods

### Animal experiments

All animal experiments were approved by the Peter MacCallum Cancer Centre Animal Experimentation Ethics Committee and conducted in congenic *Ptprca* (C57BL/6.SJL-Ptprca) mice. All mice were purchased from the Animal Resource Centre (Western Australia) and maintained under specific pathogen-free conditions. The MN and RUNX1-RUNX1T1 AML models were described previously ^14,17^. The I1DN model was generated by co-transduction of foetal liver derived HSPCs with constructs encoding *IDH1*^*R132H*^, *DNMT3A*^*R882H*^ and *Nras*^*G12D*^ (Gruber and Kats, unpublished). To generate murine leukemias, cryopreserved cells from the spleen or bone marrow of a moribund mouse were thawed and 5×10^5^ viable cells transplanted into *Ptprca* recipients. For MN and I1DN experiments, animals were pre-conditioned by sub-lethal irradiation (3.5 Gy or 5.5 Gy, respectively) administered as a single dose prior to transplant. For drug treatment, AG636 ^11^ was administered b.i.d. by oral gavage at a dose of 100mg/kg body weight. For survival studies, treatment took place on days one to five of a seven-day cycle. Doxycycline was administered *ad libitum* via doxycycline-supplemented food and water. Peripheral blood counts were performed on a Sysmex Cell Sorter (Sysmex). Mice were euthanized at pre-determined time-points or ethical end point based on clinical symptoms.

### Flow cytometry analysis and sorting

Single-cell suspensions of whole blood, bone marrow or spleen cells were incubated in ACK red cell lysis buffer (150 mM NH_4_Cl, 10 mM KHCO_3_, 0.1 mM EDTA) for 2 min and then washed in FACS buffer (PBS, 2% FBS). For blood, the lysis procedure was repeated a second time to ensure efficient removal of mature red blood cells. Cells were then re-suspended in FACS buffer and stained with fluorophore-conjugated antibodies targeted against cell surface markers (detailed list of antibodies is provided in Extended Data Table 6). Normal mouse bone marrow cells were used as a comparator and count beads were added to each sample after staining to enable quantification of cell numbers. Flow cytometric analysis was performed on FortessaX20 or Symphony flow cytometers (BD Biosciences) and data analyzed using FlowJo software (Tree Star). Cell sorting was performed a Fusion 5 flow cytometer (BD Biosciences).

### Cell culture

MN cells from the spleens of tumor bearing mice were cultured in Anne Kelso modified Dulbecco’s modified Eagle’s medium (Low glucose DMEM (Invitrogen), 4g/L glucose, 36 mg/L folic acid, 116 mg/L L-arginine HCL, 216 mg/L L-glutamine, 2g/L NaHCO_3_ and 10 mM HEPES), supplemented with 20% fetal bovine serum (FBS) (Invitrogen), 0.1 mM L-Asparagine (Sigma-Aldrich) and 1 % penicillin/streptomycin (Invitrogen). Cells were maintained at 37 °C and 10 % CO_2_. OCI-AML3, MOLM-13, were purchased from the DSMZ. HEK293T, MV4-11 and Kasumi-1 cells were purchased from ATCC. OCI-AML3, MOLM-13 and MV4-11 were cultured in RPMI-1640 (Gibco) supplemented with 20% FBS, 1 % pen/strep and 100 µM Glutamax (Invitrogen) at 37 °C and 5 % CO_2_. HEK293T cells were cultured in DMEM (Gibco) supplemented with 10% FBS and 1 % pen/strep. All cell lines in this study were routinely tested for *Mycoplasma* contamination. Cells were treated as indicated with AG636, PUGNAc (Sigma-Aldrich), AraC, Roscovitine (Assay Matrix) or DMSO (Sigma-Aldrich). For the combination therapy experiment drugs were refreshed every second day and cells passaged every 2-4. To quantify the viable cells, cells were stained with 0.2 μg/mL DAPI for 15 mins and analyzed by flow cytometry with CountBright™ Absolute Counting Beads (ThermoFisher).

### Lentiviral constructs and transduction

Cas9-mCherry (Addgene Plasmid #70182) was used to generate stable Cas9-expressing cell lines. sgRNA sequences listed in Extended Data Table 7 were cloned into the lentiGuide-Crimson backbone (Addgene Plasmid # 70683). Non-replicating lentiviruses were generated by transient co-transfection of the transfer plasmids into HEK293T together with the packaging plasmids pMDL (Addgene Plasmid #12251), pRSV-REV (Addgene Plasmid #12253) and pVSVg (Addgene Plasmid #12259). Lentivirus containing supernatants were harvested and transduced into AML cell lines with 4 μg/ml sequabrene (Sigma-Aldrich).

### Western blotting

Western blotting was performed according to standard laboratory protocols. Cells were lysed in RIPA lysis buffer (50 mM Tris-HCl pH 8, 150 mM NaCl, 1% NP-40, 0.5% sodium deoxycholate, 0.1% SDS, 0.5 U benzonase and protease inhibitors) for 30 min at 4 °C. 30µg protein lysates were resolved in 4–15% Mini-PROTEAN TGX Precast Protein Gels (Bio-Rad) and immunoblotted onto Immobilon-P PVDF membrane (Millipore). Membranes were incubated with primary and secondary antibodies and Near-Infrared Western Blot Detection performed by the Odyssey CLx imaging system (LI-COR Biosciences). The following antibodies were used: YY1 (D5D9Z) Rabbit mAb (1:1000, Cell Signaling #46395), O-GlcNAc (CTD110.6) Mouse mAb (1:1000, Cell Signaling # 9875), β-actin (1:5000, Sigma-Aldrich #A2228), IRDye® 680RD Goat anti-Mouse IgG (H + L) (1:20000-40000, LI-COR Biosciences #926-68070), IRDye® 800CW Goat anti-Rabbit IgG (H + L) (1:20000-40000, LI-COR Biosciences #926-32211).

### RNAseq and bioinformatic analysis

Sorted cKit^high^CD11b^low^ AML cells were lysed in Trizol reagent and RNA was extracted using Direct-zol RNA miniprep kit (Zymo Research) according to manufacturer’s instructions. RNA sequencing was performed at the Peter MacCallum Cancer Centre Molecular Genomics core facility. The QuantSeq 3′ mRNA-seq Library Prep Kit for Illumina (Lexogen) was used to generate libraries as per the manufacturer’s instructions. Libraries were pooled and sequenced with 75bp single end sequencing to a depth of 10 × 10^6^ reads on a NextSeq500 (Illumina). Sequencing reads were demultiplexed using bcl2fastq (v2.17.1.14) and low-quality reads Q<30 were removed. The RNA sequencing reads were trimmed at the 5’ end using cutadapt (v1.14) to remove bias introduced by random primers used in the library preparation and 3’ end trimming was performed to eliminate poly-A-tail derived reads. Reads were mapped to the reference genome (mm10) using HISAT2. Reads were counted using subread software package (v1.6.3). Differential gene expression analysis was performed using R package LIMMA (v6.5). R packages pheatmap (v1.0.12) and ggplot2 (v3.2.1) were used for figure generation. CAMERA^46^ was used for competitive testing of C2 curated gene sets from the Broad Institute’s MSigDB for mouse. GSEA (v4.0.1) was used for analyzing enrichment of gene sets and gene ontology (GO) analysis was performed using metascape (http://metascape.org).

### qPCR

RNA was extracted as outlined above. Reverse transcription was performed with the High-Capacity cDNA Reverse Transcription Kit (ThermoFisher) using 1 µg total RNA. qPCR was performed using SensiFast SYBR Hi-ROX kit (Bioline) on CFX96 Touch Real-Time PCR System (Bio-Rad). Results were analyzed using the 2^ΔΔC(t)^ method using β-actin or β-2-microglobulin for normalization. Sequences of primers used are listed in Extended Data Table 7.

### ATACseq and bioinformatic analysis

ATACseq was performed as described previously using ^47^ using sorted 5×10^4^ cKit^high^CD11b^low^ MN cells. Briefly, cells were permeabilized using 10 mM Tris-HCl pH 7.4, 10 mM NaCl, 3 mM MgCl2, 0.1% IGEPAL CA-630, 0.1% Tween-20 and tagmentation was performed using the Tagment DNA TDE1 Enzyme and Buffer kit (Illumina). Tagmented chromatin was purified using the ChIP DNA Clean & Concentrator kit (Zymo Research). Libraries were pooled and sequenced with 75bp single-end sequencing to a depth of 20 × 10^6^ reads per sample on a NextSeq500 (Illumina). Raw sequencing reads were demultiplexed using bcl2fastq (v2.17.1.14) and quality control performed with FastQC (v0.11.5). Adaptor sequences were removed using Trim Galore! (v0.4.4) and reads aligned to the mouse genome (mm10) using Bowtie2 (v2.3.3). Samtools (v1.4.1) was used for processing of SAM and BAM files including removal of duplicate reads. Peaks were called using Macs2 (v2.1.1) with the no lambda and no model settings. In R (v 4.0.2), CSAW (v1.18.0) was used to count reads in windows specified by the union of vehicle and drug treated Macs2 peaks, filter blacklisted regions (ENCODE) and perform loess normalization, then edgeR (v3.32.1) was used for differential accessibility analysis ^48^. ChIPseeker (v1.26.0) and TxDB.Mmusculus.UCSC.mm10.knownGene (v3.10.0) was used for annotation of peaks to gene features. HOMER (v4.8) was used for motif discovery. Bamtools (v2.4.1) was used to merge replicate BAM files. BAM files were converted into BigWig files using the bamCoverage function (Deeptools, v3.5.0) using the following settings (--normalizeUsing CPM --smoothLength 150 --binSize 50 -e 200). Average profile plots were generated by computing read average read density (from BigWig files) across defined genomic intervals using computeMatrix and visualised using plotProfile (Deeptools, v3.5.0).

### PSCAN analysis

PSCAN was used to identify statistically over-represented transcription factor motifs in the proximal promoters of translation genes ^30^. The analysis was performed with a window of 500 bp (−450 and +50 bp relative to TSS), using Jaspar 2018_NR TFBSs matrices. PSCAN can be accessed at: http://159.149.160.88/pscan/

### Ribosome profiling

1×10^7^ MN cells were seeded at 5×10^5^ per mL and treated with AG636 (250nM) for 24h. 1×10^7^ viable MN cells were used per biological replicate. Polysome profiling experiments were performed in cells pre-treated with 100 µg/mL cycloheximide for 5 mins. Cells were washed with ice cold PBS followed by hypotonic wash buffer (5 mM Tris-HCl (pH 7.5), 2.5mM MgCl2, 1.5 mM KCl) containing 100µg/mL cycloheximide. Cells were lysed in a hypotonic lysis buffer (5 mM Tris-HCl (pH 7.5), 2.5 mM MgCl2, 1.5 mM KCl, 100 µg/mL cycloheximide, 2mM DTT, 0.5% Triton X-100, and 0.5% sodium deoxycholate) on ice for 10 mins and lysates were pre-cleared by centrifugation to remove nuclei. The cytoplasm was collected and loaded onto a 10-40% linear sucrose density gradient (containing 20 mM Tris-HCL (pH 7.6), 100 mM KCl, 5 mM MgCl2) and centrifuged at 36,000 rpm [SW40 Ti rotor (Beckman Coulter, inc)] for 2.15h at 4°C. Gradients were fractionated (14 fractions per sample) and optical density was continuously recorded at 260nm using a sensitivity of 1 on an ISCO Tris and UA-6 UV/VIS detector (Teledyne).

### Epigenetics-targeted CRISPR screen

The screen was performed as described previously with modifications ^49^. We used a custom sgRNA library containing guides targeting 859 epigenetic regulators (four sgRNAs/gene) gene and 100 non-targeting controls (guide sequences are provided in Extended Data Table 8). To generate the library, guide sequences were PCR amplified from a CustomArray Inc oligo pool and cloned into the lentiGuide-Puro (Addgene #52963) backbone using Golden Gate cloning. For the screen, 6 × 10^6^ MN^Cas9^ cells were transduced with MOI < 0.3 to achieve single sgRNA integration per cell at an average 500-fold representation. Two days post-viral infection, transduced cells were selected by 2µg/mL Puromycin for 3 days. A time-point 0 pellet of 2 × 10^6^ live cells was harvested by centrifugation, snap frozen and stored at −80°C until required. To identify resistance or sensitization to AG636, samples were divided into 3 groups at the five days (T0) post-selection: Vehicle (1:1000 DMSO), 100nM AG636 and 250nM AG636. Drug/vehicle was refreshed every second day and cells passaged every 2-3 days, maintaining at least 500x representation by seeding 2 × 10^6^ cells in 10 mL every passage. Samples were harvested at T10 and T24 and snap frozen. Genomic DNA was extracted using the DNeasy Blood & Tissue Kit (Qiagen). Libraries were prepared by nested PCR method as described previously ^49^, pooled, purified using AMPure XP beads (Beckman Coulter) and sequenced to a depth of two million reads with single-end 75 bp sequencing on a NextSeq500 (Illumina). Sequencing reads were demultiplexed using bcl2fastq (v2.17.1.14) and low-quality reads Q < 30 were removed. The reads were trimmed using cutadapt (v1.16.5)^7^ to extract the 20 bp targeting sequence and sgRNAs that were enriched or depleted in response to AG636 relative to T0 and DMSO determined using MAGeCK algorithm (v0.5.8.1) ^37^. R packages ggplot2 (v2.2.1) and ggrepel (v0.8.0) were used for figure generation.

### Competitive Proliferation Assay

sgRNAs constructs in the lentiGuide-Crimson backbone were transduced into Cas9 expressing AML cell lines at 30-70% efficiency. Cells were treated with vehicle (1:1000 DMSO) or AG636 (250 nM or 1 µM) beginning on day 4 post transduction. Drug was refreshed every second day and cells passaged every 2-3 days. The percentage of sgRNA expressing (Crim^+^) cells were measured by flow cytometry and normalized to the initial transduction efficiency.

### Statistical Analysis

GraphPad Prism 9.0 and R version 3.6.2 software were used for statistical analysis. Statistical tests performed for each experiment are highlighted in the figure legends.

## Data and code availability

RNA-, ATAC- and CRISPR-sequencing data presented in this study has been deposited at GEO and is publicly available as of the date of publication. Deposited data is assigned under accession number - GSE181666

## Data availability

Processed and unprocessed data for RNAseq, ATACseq and CRISPR screen will be deposited to GEO and accession numbers provided prior to publication. All other data is available from the corresponding author on request.

## Acknowledgements

This work was supported by fellowships from the Victorian Cancer Agency (MCRF15003) and the Brazis family, and a research grant from the National Health and Medical Research Council of Australia (APP1099160). We thank members of the molecular genomics, animal and flow cytometry core facilities at the Peter MacCallum Cancer Centre for technical support.

## Author contributions

JS, ACL, EG and LMK performed the experiments, analyzed the data and wrote the manuscript. LMK designed the study. KS, LS, PD, LP, SJH, SJV, RWJ, GM, DBU and JM provided essential reagents and technical expertise. All authors discussed the results and approved the manuscript.

**Extended Data Fig. 1:**
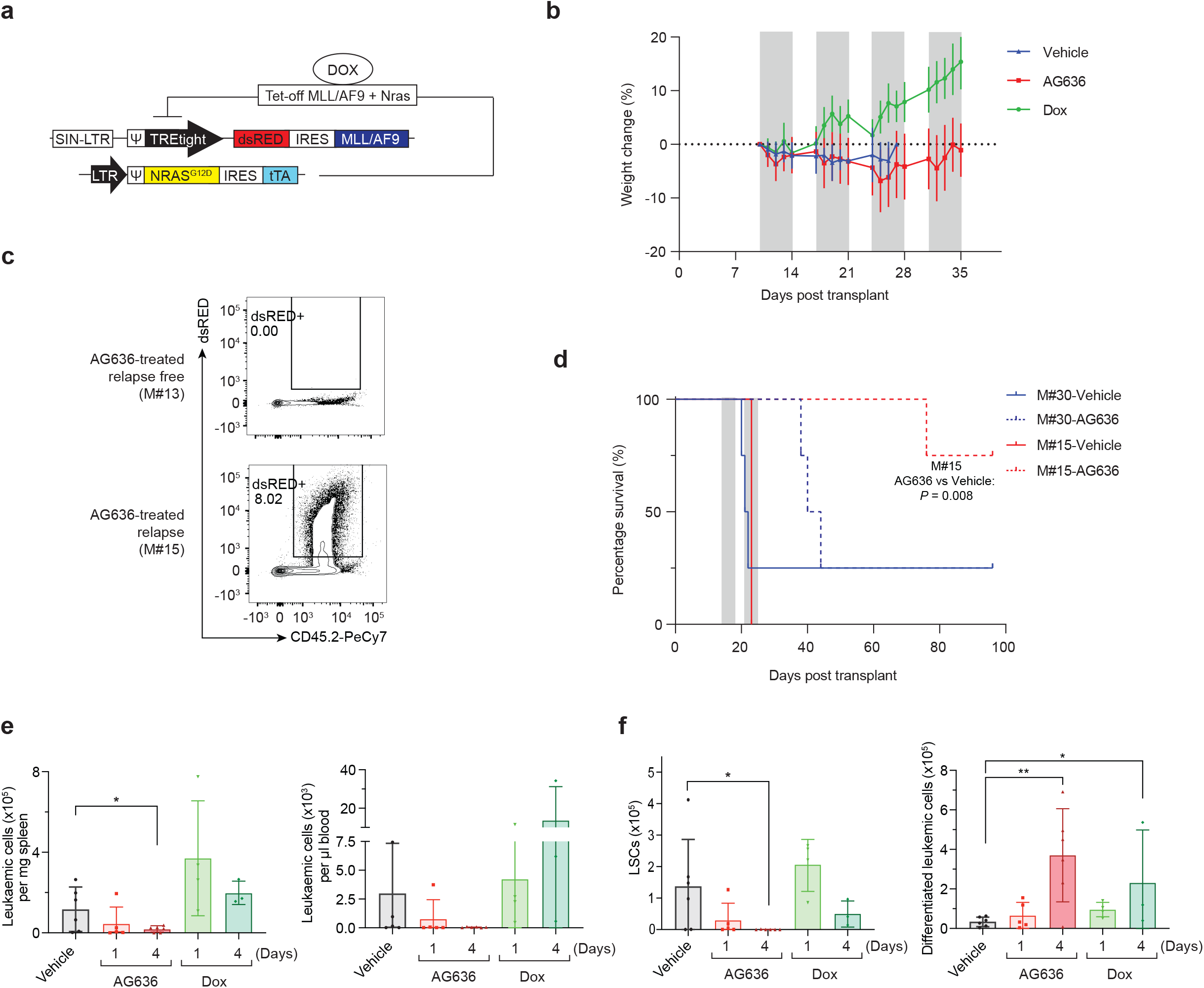
Efficacy of DHODH inhibition in the MN murine AML model. **a**, Schematic of MN model. **b**, Body weight of MN tumor-bearing mice treated with AG636. Grey bars denote treatment. **c**, Representative FACS plots of the bone marrow from a mouse with no detectable disease (M#13) and a relapsed mouse (M#15). **d**, Kaplan-Meier survival curve of secondary recipients transplanted with leukemic cells from the relapsed donor (M#15) or a control donor from the vehicle group (M#30). Grey bars denote treatment (*n* = 4 mice/group, median survival is 21.5 for vehicle-treated M#30, 42 for AG636-treated M#30, 23 for vehicle-treated M#15 and not reached for AG636-treated M#15, *P* value was calculated by Log-rank test). **e**, number of MN cells in the spleen and peripheral blood quantified by flow cytometry. **f**, number of LSCs (CD11b^low^cKit^high^FcgR^+^) and differentiated leukemic cells (CD182^+^Ly6G^+^) in the bone marrow (*n =* 3-6 mice/group). Data in **e-f** are presented as mean ± s.d.; *P* values were calculated using a one-tailed Student’s unpaired t-test. \**P* < 0.05, \*\**P* < 0.01, \*\*\**P* < 0.001; Dox - doxycycline.

**Extended Data Fig. 2:**
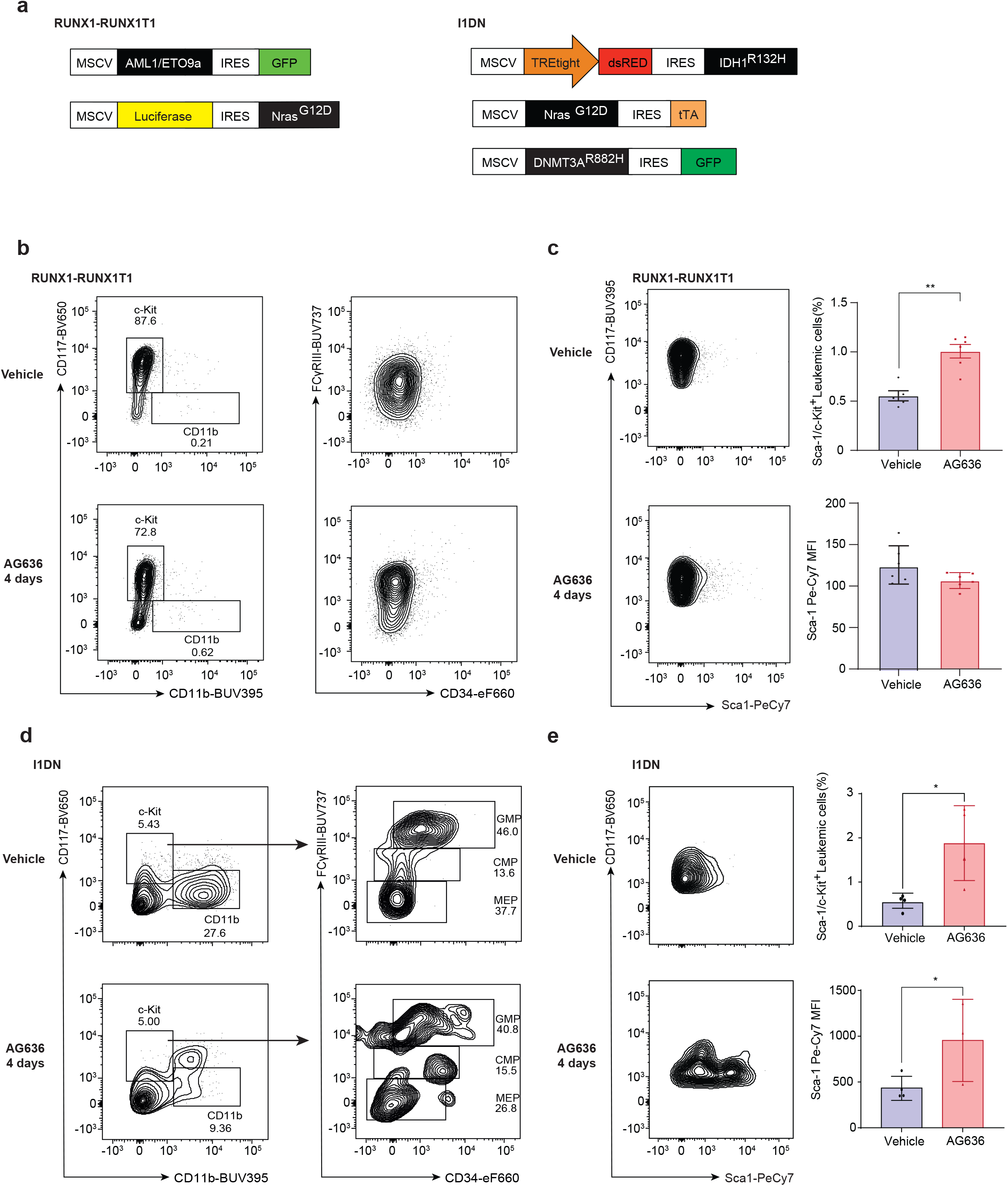
Efficacy of DHODH inhibition in the RUNX1-RUNX1T1 and I1DN murine AML models. **a**, Schematic of RUNX1-RUNX1T1 and I1DN models. **b-e**, Representative FACS plot of the bone marrow of RUNX1-RUNX1T1 (**b-c**) or I1DN (**d-e**) tumor bearing mice showing expression of the indicated cell surface markers. Quantification of Sca1 expression is shown. Data are presented as mean ± s.d.; *P* values were calculated using a one-tailed Student’s unpaired t-test. \**P* < 0.05.

**Extended Data Fig. 3:**
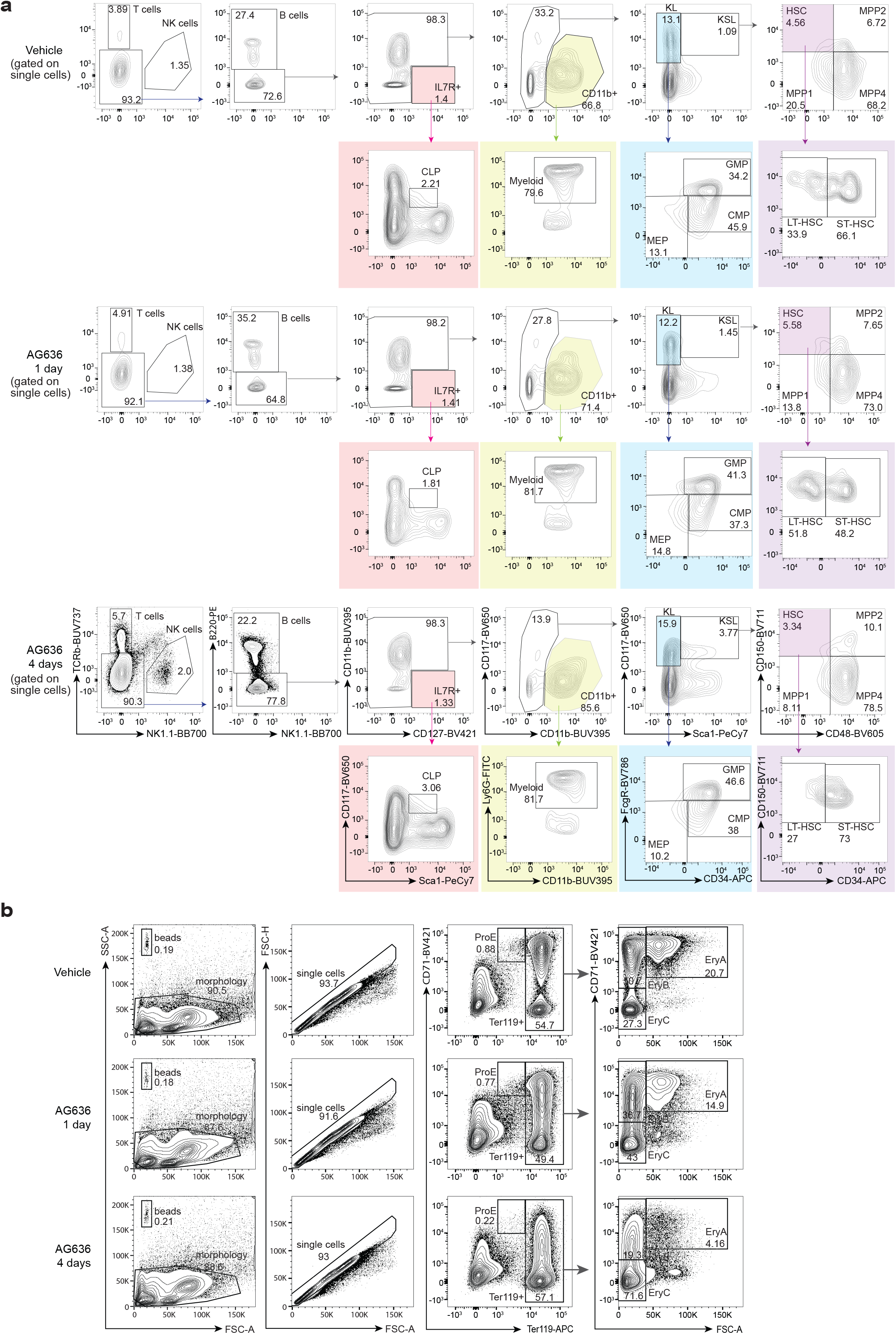
Identification of bone marrow sub-population by FACS. **a**, FACS gating strategy for bone marrow mononuclear cells (MNCs). **b**, FACS gating strategy for erythroid cells.

**Extended Data Fig. 4:**
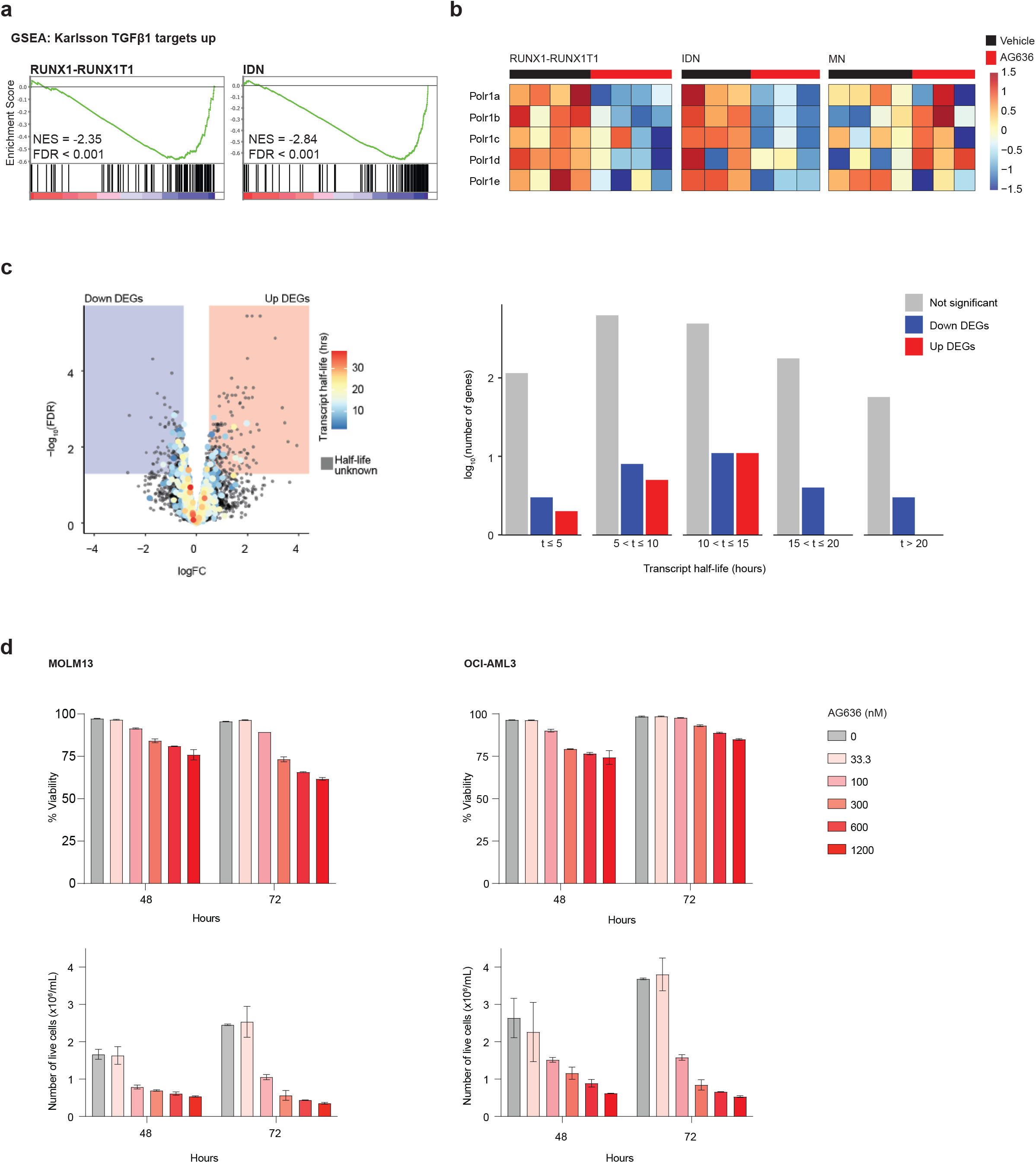
Transcriptional effects of DHODHi in AML. **a**, Bar code plots showing downregulation of TGF-β signaling in RUNX1-RUNX1T1 and I1DN murine AML models following AG636 treatment. **b**, Gene expression heat map showing down-regulation of genes encoding components of RNA polymerase I in RUNX1-RUNX1T1 and I1DN murine AML models following AG636 treatment. **c**, Volcano plot of gene expression in MN cells, highlighting the average transcript half-life of each gene (left) and bar chart of the number of genes with transcript half-lives in the given interval for significant DEGs (right). Transcript half-life was provided by ^28^. **d**, Cell viability of human AML cell lines treated with AG636 (*n =* 2 biological replicates; data represented as mean ± s.e.m).

**Extended Data Fig. 5:**
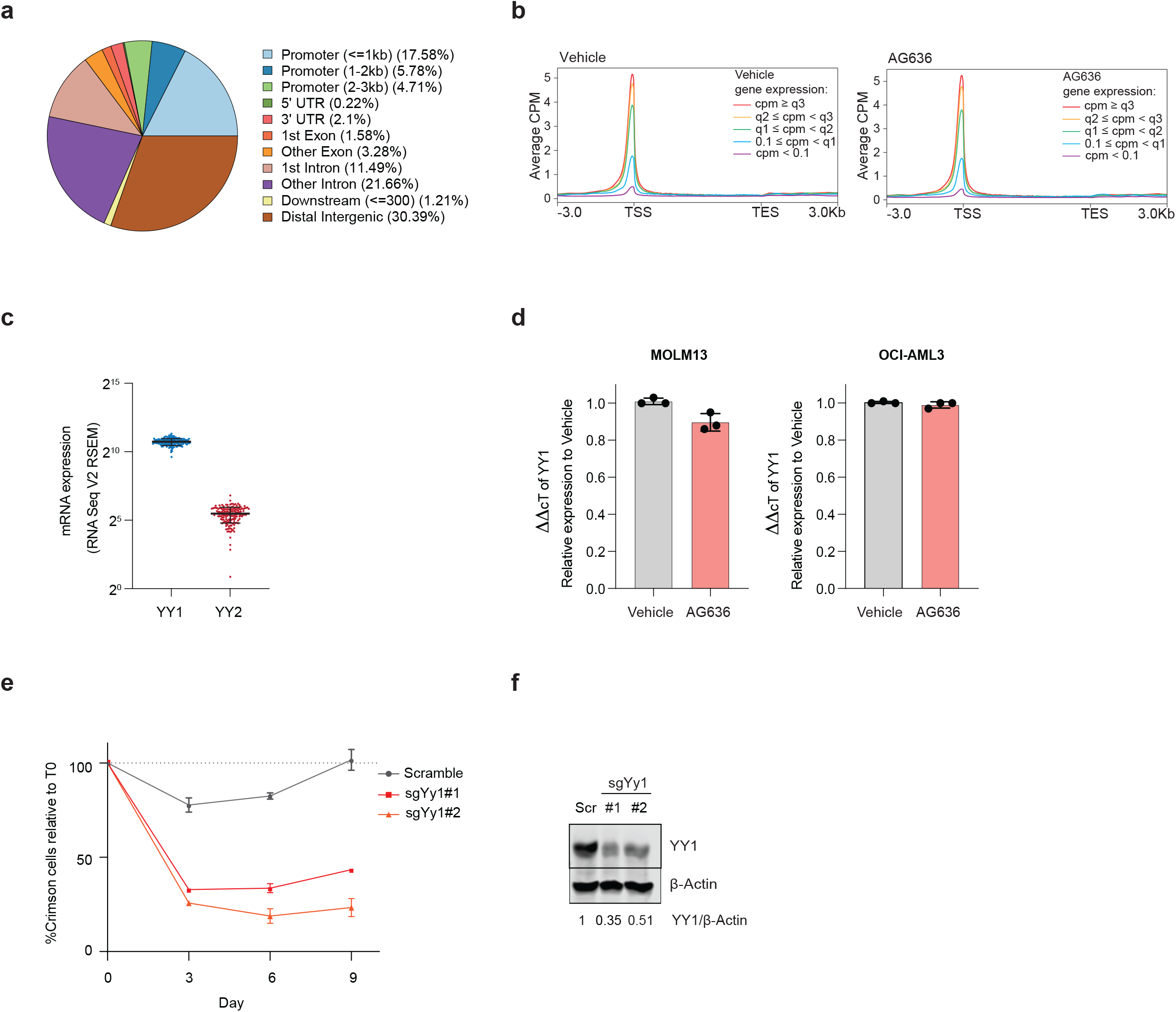
YY1 is a downstream target of AG636 in AML. **a-b**, ATAC sequencing performed on cKit^high^CD11b^low^ MN cells sorted from mice treated with AG636 or vehicle for 2 days (*n =* 3 mice/group). **a**, Pie chart showing association between regions of accessible chromatin and different genome features in untreated MN cells. **b**, Metaplot of ATACseq signal at genes grouped by gene expression. Quantile values were calculated on genes whose expression was ≥ 0.1 CPM. **c**, Expression of YY1 and YY2 in AML patient samples in the TCGA database ^31^ (data represented as mean ± s.d.). **d**, qPCR of *YY1* expression in human AML cell lines treated with AG636 for 24 hours (*n =* 2 biological replicates; data represented as mean ± s.e.m). **e**, Proliferative competition assays in MN cells transduced with two independent YY1-targeting sgRNAs or a non-targeting control sgRNA (*n =* 2 biological replicates; data represented as mean ± s.e.m). **f**, Western blot of YY1 expression in MN cells transduced with two independent YY1-targeting sgRNAs or a non-targeting control sgRNA.

**Extended Data Fig. 6:**
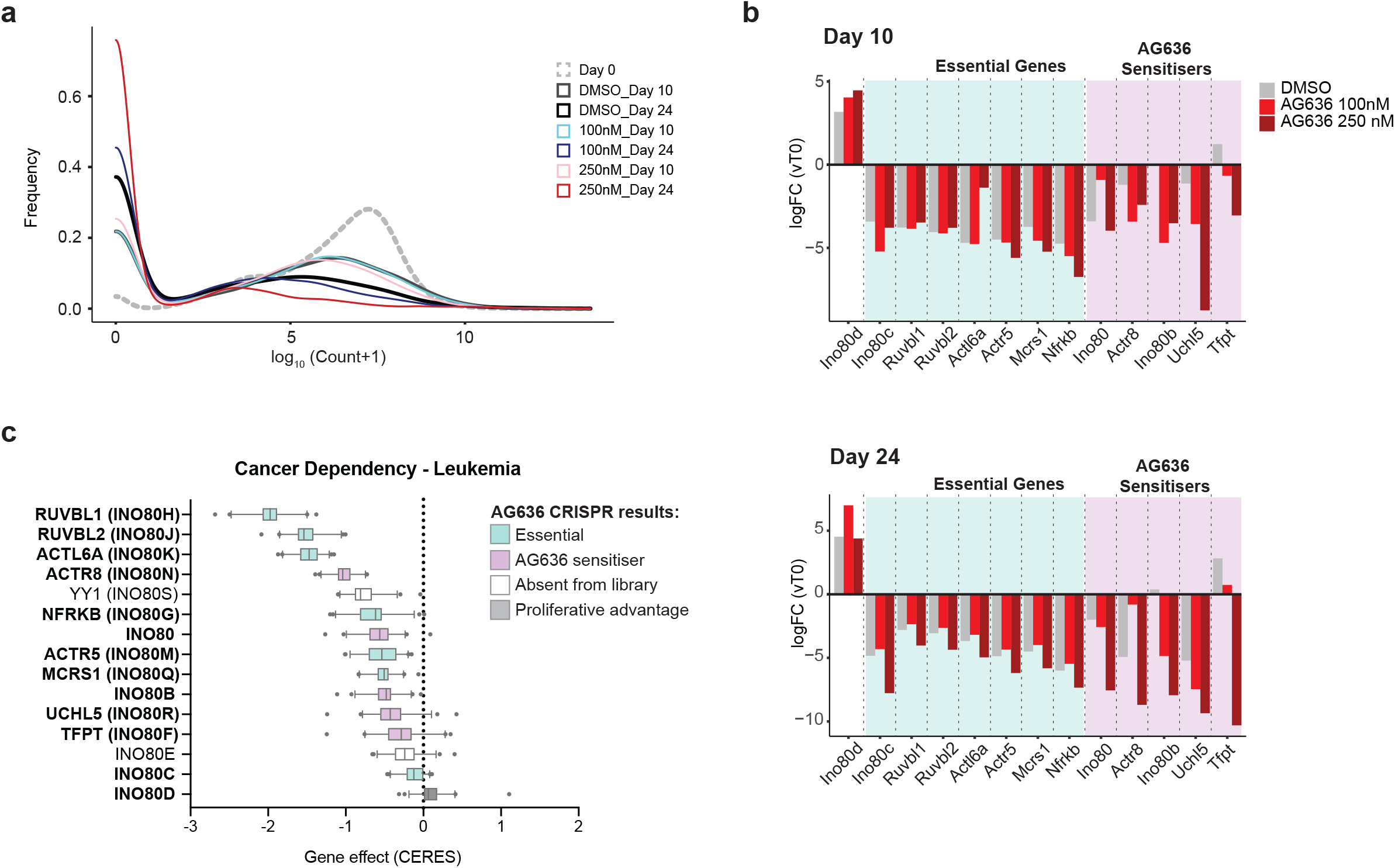
Identification of genes that increase or decrease sensitivity of AML cells to DHODHi. **a**, Distribution of sgRNA counts in the various conditions during the CRISPR screen. **b**, Average fold change in sgRNA counts compared with time-point 0 for all sgRNAs targeting components of the INO80 complex in the various conditions in the CRISPR screen. **c**, Gene dependencies in AML cell lines from the DepMap database ^35^, Centre line; median; box limits, from the 25^th^ to 75^th^ percentiles; whiskers, from the 5^th^ to 95^th^ percentiles.

**Extended Data Fig. 7:**
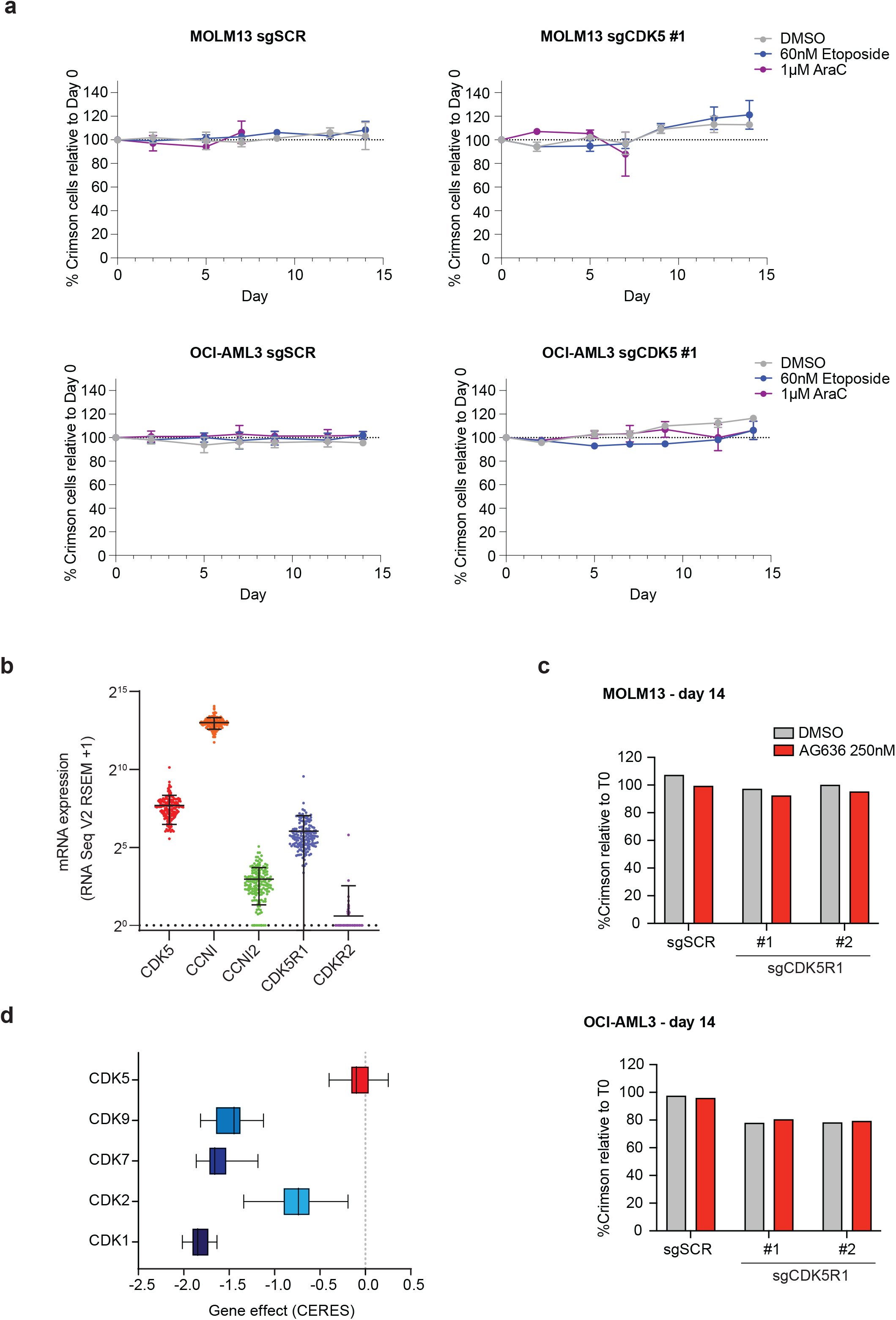
CDK5/CCNI expression affects response to DHODHi in AML. **a**, Proliferative competition assays in human AML cell lines transduced with CDK5 targeting sgRNAs and cultured in AraC (*n =* 2 biological replicates; data represented as mean ± s.e.m). **b**, Expression of *CDK5* and its activators in AML patient samples in the TCGA database ^31^ (data represented as mean ± s.d.). **c**, Proliferative competition assays in human AML cell lines transduced with indicated sgRNAs and cultured in AG636 or DMSO. **d**, Gene dependencies in AML cell lines from the DepMap database ^35^, Centre line; median; box limits, from the 25^th^ to 75^th^ percentiles; whiskers, from the 5^th^ to 95^th^ percentiles..

**Extended Data Table 1: Cell surface markers used to identify bone marrow sub-populations**.

**Extended Data Table 2: Differential gene expression in cKit**^**high**^**CD11b**^**low**^ **MN cells sorted from AG636, doxycycline or vehicle treated mice**.

**Extended Data Table 3: Differential gene expression in cKit**^**high**^**CD11b**^**low**^ **RUNX1-RUNX1T1 AML cells sorted from mice treated with AG636 or vehicle for four days**.

**Extended Data Table 4: Differential gene expression in cKit**^**high**^**CD11b**^**low**^ **I1DN AML cells sorted from mice treated with AG636 or vehicle for four days**.

**Extended Data Table 5: Pscan analysis of transcription factor motifs enriched within the proximal promoters of translation genes**.

**Extended Data Table 6: Antibodies used in this study**

**Extended Data Table 7: Primer and sgRNA sequences**

**Extended Data Table 8: Epigenetics-targeted CRISPR library guides sequences**

